# Single-cell multiomics sequencing reveals the functional regulatory landscape of early embryos

**DOI:** 10.1101/803890

**Authors:** Yang Wang, Peng Yuan, Zhiqiang Yan, Ming Yang, Ying Huo, Yanli Nie, Xiaohui Zhu, Liying Yan, Jie Qiao

## Abstract

Extensive epigenetic reprogramming occurs during preimplantation embryo development and is accompanied by zygotic genome activation (ZGA) and first cell fate specification. Recent studies using single-cell epigenome sequencing techniques have provided global views of the dynamics of different epigenetic layers during this period. However, it remains largely unclear how the drastic epigenetic reprogramming contributes to transcriptional regulatory network. Here, we developed a single-cell multiomics sequencing technology (scNOMeRe-seq) that enables profiling of genome-wide chromatin accessibility, DNA methylation and RNA expression in the same individual cell with improved performance compared to that of earlier techniques. We applied this method to analyze the global dynamics of different molecular layers and their associations in mouse preimplantation embryos. We found that global DNA methylation remodeling facilitates the reconstruction of genetic lineages in early embryos and revealed that the gradual increases in heterogeneity among blastomeres are driven by asymmetric cleavage. Allele-specific DNA methylation pattern is maintained throughout preimplantation development and is accompanied by allele-specific associations between DNA methylation and gene expression in the gene body that are inherited from oocytes and sperm. Through integrated analyses of the collective dynamics between gene expression and chromatin accessibility, we constructed a ZGA-associated regulatory network and revealed coordination among multiple epigenetic layers, transcription factors (TFs) and repeat elements that instruct the proper ZGA process. Moreover, we found that inner cell mass (ICM)/trophectoderm (TE) lineage-associated cis-regulatory elements are stepwise activated in blastomeres during post-ZGA embryo stages. TE lineage-specific TFs play dual roles in promoting the TE program while repressing the ICM program, thereby separating the TE lineage from the ICM lineage. Taken together, our findings not only depict the first single-cell triple-omics map of chromatin accessibility, DNA methylation and RNA expression during mouse preimplantation development but also enhance the fundamental understanding of epigenetic regulation in early embryos.

## Introduction

In mammals, embryo development starts from a unified zygote. Coincident with the first several zygotic cleavages, early embryos activate the zygotic genome, restore totipotency and further generate the inner cell mass (ICM) and trophectoderm (TE) during preimplantation development^1–3^. Failures in zygotic genome activation (ZGA) or ICM/TE lineage specification can cause early embryo developmental arrest and implantation failure in both mice and humans. With advances in low-input and single-cell epigenome sequencing, recent studies have revealed that extensive global epigenetic reprogramming, for instance, reprogramming of DNA methylation (Met), chromatin accessibility (Acc) and histone modifications, occurs in early embryos during this period^4–14^. However, it remains to be explored how these epigenome reconfigurations contribute to the establishment of proper regulatory networks of early embryos.

Acc is a hallmark of cis-regulatory elements (CREs), such as promoters and enhancers, that act coordinately with transcription factors (TFs) and epigenetic modifications to finely regulate the transcriptional activity of downstream genes and establish cell-type specific regulatory networks^15, 16^. The currently optimized low-input open-chromatin sequencing techniques, such as the assay for transposase-accessible chromatin using sequencing (ATAC-seq) and low-input DNase I sequencing (liDNase-seq), are able to detect genome-wide dynamics of Acc in early embryos. However, the signal of the open regions reflects the average signal of the mixed sample, which may be confounded by highly heterogeneous and asynchronized blastomeres and even abnormal embryos^6, 10, 17, 18^. Single-cell ATAC-seq detects only thousands of informative reads per cell on average, which might limit its application with the scarce resources of early embryos^19–21^. Moreover, because of the high heterogeneity of early blastomeres, a functional understanding of epigenomic changes requires knowledge of the transcriptional output from one individual cell. Recently, different single-cell epigenome sequencing methods and transcriptome sequencing methods have been combined to profile different combinations of molecular layers from the same individual cell, providing opportunities to explore the associations between different molecular layers^22–26^. However, factors compromising the quality of data from current single cell multi-omics technologies, such as poor genome coverage or low gene number detection, might constrain the precise interpretation of the associations between different molecular layers.

Here, we describe a technique called **s**ingle-**c**ell **n**ucleosome **o**ccupancy, **me**thylome and **R**NA **e**xpression **seq**uencing (scNOMeRe-seq) that effectively combines single-cell nucleosome occupancy and methylome sequencing (scNOMe-seq) with Multiple Annealing and dC-Tailing-based Quantitative single-cell RNA sequencing (MATQ-seq), showing improved performance for profiling of multiple molecular layers from the same individual cell^4, 27, 28^. We applied scNOMeRe-seq to analyze genome-wide Acc, Met and RNA expression (Expr) in mouse preimplantation embryos at single-cell resolution and to provide a comprehensive overview of the functional regulatory landscape in early embryos.

## RESULTS

### scNOMeRe-seq profiles in mouse preimplantation embryos

To simultaneously detect genome-wide Acc, Met and Expr in the same individual cell, we developed a single-cell multiomic sequencing method, scNOMeRe-seq, by combining scNOMe-seq and MATQ-seq (Fig. 1a). We employed this method to profile 233 single cells isolated from mouse preimplantation embryos at different stages with RNA data from 221 (94.8%) single cells and DNA data from 218 (93.4%) single cells passed our stringent criteria, showing a high success rate (Extended Data Fig. 1a, f and g; Supplementary Table 1). The DNA data showed better genome coverages than those in a previous study using single-cell chromatin overall omic-scale landscape sequencing (scCOOL-seq) (this study: WCG 3.49 million, 15.8%, GCH 31.0 million, 15.5% on average per cell; scCOOL-seq: WCG 2.24 million, 10.1%, GCH 19.7 million, 9.8% on average per cell)^4^. The RNA dataset showed high accuracy, high reproducibility, even coverage through genic regions, and high detection sensitivity for genes expressed at low levels (Extended Data Fig. 1b-e)^29^. More importantly, our RNA dataset could faithfully distinguish between ICM and TE cells in embryonic day (E) 3.5 blastocysts (Extended Data Fig. 1h-i), further confirming the high quality of our RNA data obtained from scNOMeRe-seq.

**Figure 1.**
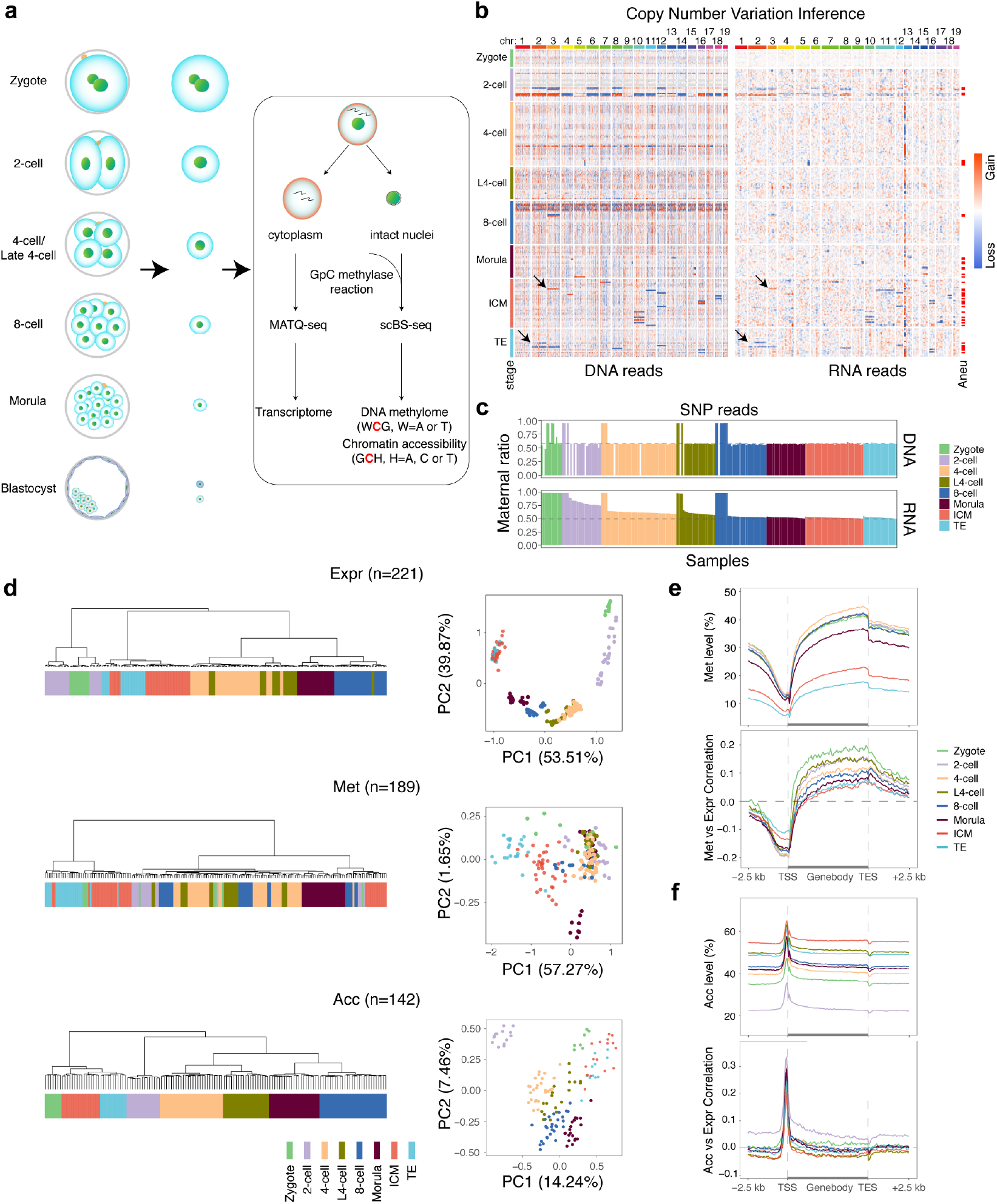
scNOMeRe-seq profiles in mouse preimplantation embryos. (**a**) Schematic illustration of scNOMeRe-seq, including key steps, methods of library preparation and mouse preimplantation stages analyzed in this work outlined in the text. (**b**) Heat map showing copy number variations (CNV) inferred by DNA reads (left) and gene expression level (right, normalized each gene by average closest 100 genes) in mouse preimplantation blastomeres. Arrows indicate the examples of matched CNV inferences from DNA and RNA reads. (**c**) Bar plot showing the ratios of SNP tracked maternal DNA (top) or RNA (bottom) reads in total SNP tracked parental reads in each individual cell across preimplantation development. (**d**) Unsupervised clustering (left) and principle component analysis (right) of preimplantation blastomeres using gene expression level (top), DNA methylation level of 5 kilobases (kb) tiles (middle) and chromatin accessibility of all stage merged NDRs (bottom). n, the cell numbers of each dataset. (**e**) Profiles showing DNA methylation level (top), the weighted Pearson correlation coefficients of DNA methylation level vs gene expression level (bottom) along the gene bodies and 2.5 kb upstream of the transcription start sites (TSS) and 2.5 kb downstream of the transcription end sites (TES) of all genes for each stage. Met, WCG methylation; Expr, RNA expression. (**f**) Profiles showing chromatin accessibility (top), the weighted Pearson correlation coefficients of chromatin accessibility vs gene expression level (bottom), along the gene bodies and 2.5 kb upstream of the TSS and 2.5 kb downstream of the TES of all genes for each stage. Acc, GCH methylation.

To detect the abnormal blastomeres in our early embryos, we analyzed the copy number variations (CNVs) with the RNA data and DNA data for each individual cell. Consistent results were obtained from both datasets regarding the inferred CNVs, even for the partial chromosome CNVs (Fig. 1b and Extended Data Fig. 2a). Unexpectedly, we also found several parthenogenetic (PG) embryos among our detected early embryos using single-nucleotide polymorphism (SNP)-separated allelic reads (Fig. 1c and Extended Data Fig. 2a). Then, we sought to determine whether the embryo abnormalities would cause aberrant embryo development. Notably, along the developmental trajectory inferred from the RNA dataset, most PG blastomeres showed delayed development after ZGA compared to that of normal and aneuploid blastomeres (Extended Data Fig. 2b-c). Although only 58% of ZGA genes were activated in PG blastomeres at the 2-cell stage, the PG blastomeres became more similar to the rest of the blastomeres beginning at the same stage after ZGA, indicating that the PG embryos were able to go through at least partial ZGA and develop to further stages (Extended Data Fig. 2b and d). The Met in PG blastomeres were clearly distinct from that in normal and aneuploid blastomeres, while Acc did not differ among PG, normal and aneuploid blastomeres (Extended Data Fig. 2f-g and 3a). Together, these results revealed that the aneuploid cells were able to undergo preimplantation development as well as proper epigenetic reprogramming; however, the PG cells showed delayed development and an aberrant Met pattern.

Furthermore, k-means clustering with the Acc of the transcription start site (TSS) could not distinguish the abnormal cells from the normal cells; however, the cells from the two clusters showed significant differences in global Acc levels and correlations between Acc and Expr at the TSS regions for each stage (Extended Data Fig. 3b). Notably, the cells from the cluster with the relatively higher Acc level (cluster_2) consistently showed lower correlations between Acc and Expr at the TSS regions, without differences in global Met levels, and correlations between Met and Expr were observed between the two clusters (Extended Data Fig. 3b). A previous study revealed globally increased Acc at S phase during mitosis; such changes in Acc are not associated with transcriptional regulation but are potentially linked to DNA duplication^30^. Therefore, these two clusters in each stage reflected the highly asynchronous cell cycles among the blastomeres of the early embryos. Furthermore, we detected the nucleosome-depleted regions (NDRs) using an aggregated Acc dataset from single cells in each cluster at each stage. Regardless of the genome coverage, cluster_1 (low Acc level and high correlation between Acc and Expr) exhibited more NDRs than cluster_2 for each stage (Extended Data Fig. 3c-d). The NDRs in cluster_1 at each stage showed greater fractions overlapping with previously defined open chromatin in early embryos than those in cluster_2 (Extended Data Fig. 3e-f)^4, 6, 10^. Together, these results reveal that the heterogeneity of Acc among blastomeres of the same stage might mainly derive from highly asynchronous blastomere cell cycles and suggest that the mixed asynchronous populations may increase the background noise and compromise the discovery of transcription-related Acc signals.

To focus on transcriptional regulation-related epigenome characteristics during preimplantation development, the Acc datasets of cells from cluster_2 and the Met datasets of PG cells were removed for downstream analysis. Then, we explored the dynamics and associations of different molecular layers in each single cell during preimplantation development. Both unsupervised clustering and principal component analysis (PCA) revealed that cells of the same stage clustered more closely within each molecular layer, consistent with the findings of previous studie (Fig. 1d)^4^. The global Met levels were relatively stable in earlier stages but sharply decreased at the blastocyst stage (Fig. 1e, Extended Data Fig. 2e). The correlations between Met and Expr in the TSS and gene body regions showed the highest associations in zygotes and gradually decreased in the following stages (Fig. 1e). In contrast, global Acc was drastically decreased at the 2-cell stage and restored at the 4-cell stage before gradually increasing at later stages (Fig. 1f, Extended Data Fig. 2e). Notably, the correlations between Acc and Expr at the TSS regions were the most positive at the 2-cell stage among all preimplantation stages, coinciding with ZGA; this finding suggests that the drastic Acc reprogramming at the 2-cell stage might contribute to proper ZGA (Fig. 1f).

### Reconstruction of genetic lineages reveals the source of heterogeneity in early embryos

Given the insufficient maintenance of Met levels during mitosis in early embryos, a previous study sought to reconstruct the genetic lineages of 4-cell embryos using single-cell genome-wide CpG Met datasets in both humans and mice and successfully elucidated the lineages^5^. To test whether our single-cell Met (WCG) datasets could be used to infer the genetic lineages of early embryos, we first computed the pairwise correlations among blastomeres in each individual 4-cell or late 4-cell embryo (see Methods). We repeatedly observed two pairs of cells with highly negatively correlated Met levels in each individual embryo, consistent with previous findings (Fig. 2a, b and d)^5^. Then, we validated that the two cells in each pair originated from the same mother 2-cell blastomere (Fig. 2e). Interestingly, we also observed a conserved pairwise correlation of Met levels among blastomeres for each analyzed 8-cell embryo, implying that it might be possible to reconstruct genetic lineages for 8-cell embryos using single-cell Met datasets (Fig. 2c and d). To verify the Met correlation patterns among blastomeres at the 8-cell stage derived from the same blastomeres in 2-cell and 4-cell embryos, we microinjected FITC to label one blastomere each in 2-cell and 4-cell embryos and performed single-cell bisulfite sequencing (scBS-seq) for each individual cell when these embryos developed to the 8-cell stage. Blastomeres in 8-cell embryos derived from the same blastomeres in 4-cell embryos exhibited highly positively correlated Met levels; in contrast, 2 pairs of blastomeres in 8-cell embryos derived from the same blastomeres in 2-cell embryos exhibited highly positively correlated Met levels within pairs, but cells from different pairs exhibited highly negatively correlated Met levels (Fig. 2f-g). Therefore, these results demonstrate that we can accurately construct the full lineages from the zygote stage to the 8-cell embryo stage using single-cell Met datasets.

**Figure 2.**
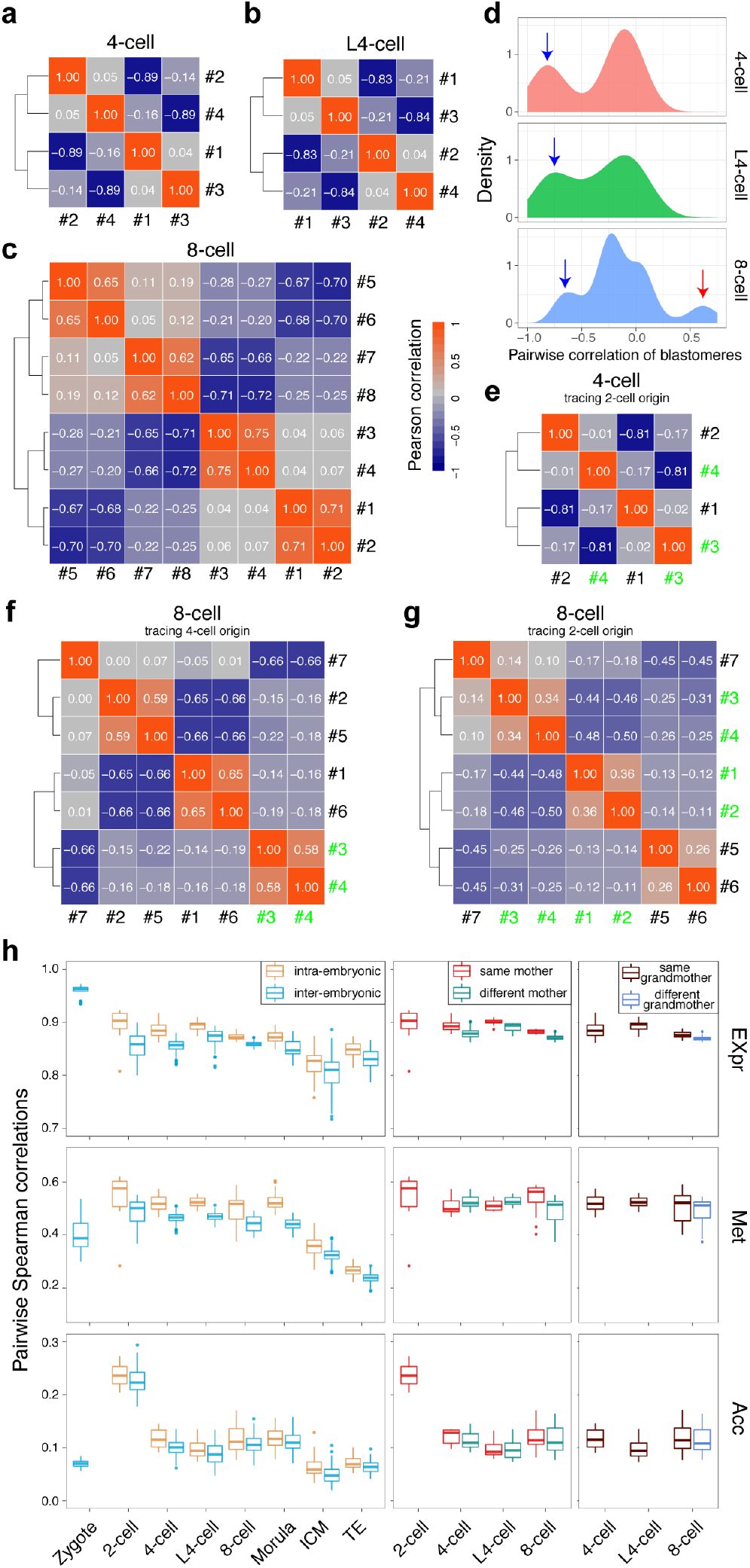
Reconstruction of genetic lineages reveals the source of heterogeneity in early embryos. (**a-c**) Heat map showing the Pearson correlation coefficients in representative 4-cell embryo (**a**), late 4-cell embryo (**b**), and 8-cell embryo (**c**), using z-score normalized DNA methylation level of 1 millionbases (Mb) bins in individual blastomere from the same embryo. The numbers in white color showing the Pearson correlation coefficients. (**d**) Distribution of the pairwise Pearson correlation coefficients for DNA methylation level of 1 Mb bins in individual blastomere from the same 4-cell embryo (top), late 4-cell embryo (middle) and 8-cell embryo (bottom). Blue arrows indicate the pairs of blastomeres from the same 2-cell blastomere, and the red arrow indicates the pairs of blastomeres from the same 4-cell blastomere. (**e**) Heat map showing the Pearson correlation coefficients in a 4-cell embryo. #3-4 (green labeled) cells divided from the same 2-cell blastomere validated by injection with FITC-dye. The numbers in white color showing the Pearson correlation coefficients. (**f**) Heat map showing the Pearson correlation coefficients in an 8-cell embryo. #3-4 (green labeled) cells divided from the same 4-cell blastomere validated by injection with FITC-dye. The numbers in white color showing the Pearson correlation coefficients. (**g**) Heat map showing the Pearson correlation coefficients in an 8-cell embryo. #1-4 (green labeled) cells divided from the same 2-cell blastomere validated by injection with FITC-dye. The numbers in white color showing the Pearson correlation coefficients. (**h**) Box plot showing the pairwise Spearman correlation coefficients of RNA expression level (top), DNA methylation level of 5 kb bins (middle) and chromatin accessibility of NDRs (bottom) at indicated stages. intra-embryonic, within the same embryo; inter-embryonic, in different embryos of the same stage; same mother, the blastomeres origin from a same mother blastomere; same grandmother, the blastomeres origin from a same grandmother blastomere.

Furthermore, we investigated when unified zygotes generate heterogeneity in different molecular layers among blastomeres. We first computed the correlations between blastomeres within each embryo (intraembryonic correlations) versus those between blastomeres from different embryos (interembryonic correlations) at the same stage for each molecular layer. We found that the intraembryonic correlations were consistently higher than the interembryonic correlations for each molecular layer throughout the preimplantation development stages, suggesting highly asynchronous development among different embryos at the same stage (Fig. 2h). Moreover, the correlations in Expr levels were highest at the zygote stage and gradually decreased at later stages, suggesting that the heterogeneity among blastomeres in the same embryo was generated during ZGA and gradually increased with preimplantation development (Fig. 2h). We also noticed that the correlations in both the Met and Acc levels were highest at the 2-cell stage, indicating that the epigenome was robustly reprogrammed for each individual cell during ZGA (Fig. 2h). Leveraging this lineage tracing information, we further explored the dynamics of heterogeneity between daughter cells during the first three cleavages. In the transcriptome, the correlations between blastomeres from the same mother cells gradually decreased during the first three cleavages, whereas the correlations between blastomeres from the same mother cells at the late 4-cell stage were comparable to those at the 4-cell stage (Fig. 2h). Moreover, the correlations between blastomeres from the same grandmother cells were higher than those of blastomeres from different grandmother cells in 8-cell embryos (Fig. 2h). These results demonstrated that asymmetric cleavage might have been the major source of the transcriptome heterogeneity. Although the heterogeneity in the epigenome seemed not to be associated with asymmetric cleavage, we notably observed that the correlations in Met levels between blastomeres from the same mother cells were higher in 8-cell embryos than in 4-cell and late 4-cell embryos, indicating that increased Met maintenance occurs to some extent during DNA duplication at the 4-cell stage (Fig. 2h).

### Allele-specific regulation of gene expression in early embryos

Drastic epigenetic reprogramming occurs in parental genomes after fertilization. The Acc levels of the parental genomes were comparable in most individual cells throughout preimplantation development (Fig. 3a). The Met level in the paternal allele was consistently higher than that in the maternal allele for each individual zygote (Fig. 3b). From the 2-cell to the 8-cell stage, the global differences in Met levels between parental alleles varied in the different individual cells; however, after the morula stage, the Met level in the maternal allele was consistently higher than that in the paternal allele for each individual cell (Fig. 3b). In addition, we found that 16.7%-29.7% of regions showed significant allelic differences (FDR < 0.01) in Acc levels, and 7.9%-40.2% of regions showed significant allelic differences (FDR < 0.01) in Met levels across preimplantation stages (Extended Data Fig. 4a-b). The differences in allelic Acc levels were widely distributed in the whole genome and showed no preference for a particular parental allele (Fig. 3c, Extended Data Fig. 4c). Notably, the maternal hypermethylated regions were highly enriched in genic regions, whereas the paternal hypermethylated regions were highly enriched in distal intergenic regions throughout the preimplantation stages, consistent with previous findings (Fig. 3d, Extended Data Fig. 4d)^4^. Given that the oocyte genome is highly methylated at actively transcribed genic regions and hypomethylated at intergenic regions, while the sperm genome is highly methylated at intergenic regions, our results indicate that global differences in Met levels between parental alleles in gametes could be largely maintained throughout preimplantation development^4, 12^.

**Figure 3.**
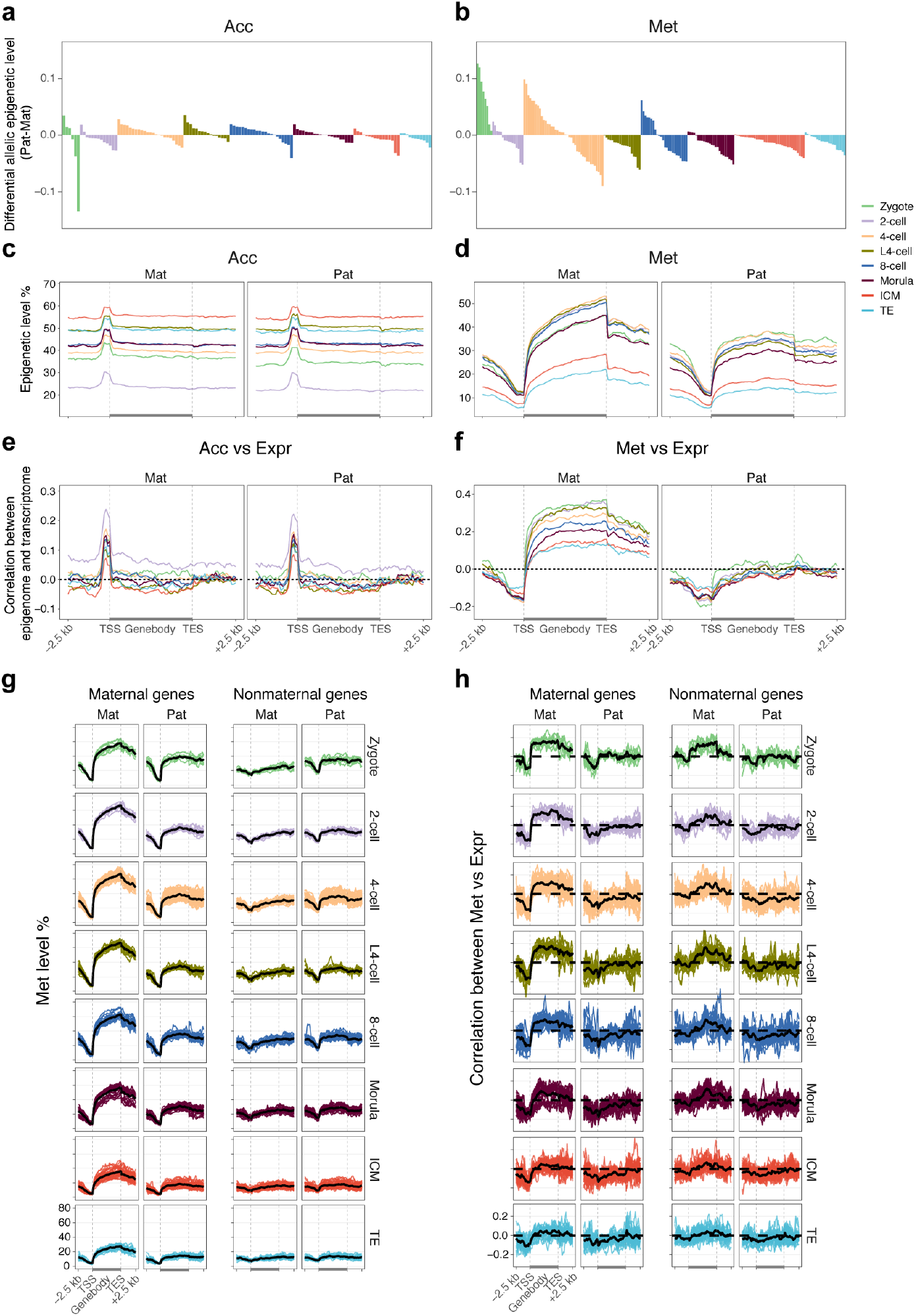
Allele-specific regulation of gene expression in early embryos. (**a-b**) Bar plot showing differences of global chromatin accessibility (**a**) and DNA methylation level (**b**) between paternal and maternal genomes in each individual cell of preimplantation embryos. (**c-d**) Allelic chromatin accessibility (**c**) and DNA methylation level (**d**) around genic regions of each stage. Mat, maternal allele; Pat, paternal allele. (**e**) The Pearson correlations between chromatin accessibility around genic regions of maternal allele or paternal allele vs gene expression level of each stage. (**f**) The Pearson correlations between DNA methylation level around genic regions of maternal allele or paternal allele vs gene expression level of each stage. (**g**) Allelic DNA methylation level around genic regions of maternal (TPM ≥ 1 in zygote stage) and nonmaternal genes in each stage. (**h**) The Pearson correlations between DNA methylation level around genic regions of maternal allele or paternal allele vs expression level of maternal and nonmaternal genes.

We next sought to determine whether the allelic epigenome differences were associated with allelic transcriptional regulation. First, we overlapped the differential allelic epigenetic regions with known imprinting control regions (ICRs)^31^. Four germline ICRs overlapped with our differential allelic Acc regions, and all showed corresponding differential allelic Acc patterns in at least one preimplantation stage (Extended Data Fig. 4e). In the other hand, six known germline ICRs overlapping with differential allelic Met regions showed the expected differential allelic Met patterns throughout preimplantation development, validating the accuracy of our analysis (Extended Data Fig. 4f). Furthermore, we assessed the correlations between allelic epigenetic modification levels and Expr in each individual cell. Both parental alleles showed similar correlation patterns between allelic Acc and Expr, mimicking the overall Acc vs Expr associations (Fig. 3e). Notably, we observed clearly different correlation patterns between allelic Met and Expr in parental alleles: the Met levels of the paternal genome at the gene body regions showed no correlations with Expr, unlike those in the maternal genome (Fig. 3f). To determine whether the allele-specific correlations between Met and Expr at gene bodies were caused by inherent correlations from maternal factors, we further compared the correlations between allelic Met and the Expr of maternal genes (transcript per million mapped reads (TPM) ≥ 1 in the zygote) with nonmaternal genes. The Met levels at gene body regions were clearly higher for maternal genes than for nonmaternal genes in maternal alleles throughout preimplantation stages (Fig. 3g). As expected, major Met differences between parental alleles were observed in maternal genes but not in nonmaternal genes (Fig. 3g). Furthermore, we found that the correlations between maternal Met and Expr at gene body regions were clearly weaker in nonmaternal genes than in maternal genes, indicating that the observed positive correlations between maternal Met and Expr at gene body regions were mainly inherited from oocytes (Fig. 3h).

### scNOMeRe-seq reveals a ZGA-associated regulome

To reveal ZGA-associated CREs, we measured the correlations between the Acc of each promoter/distal NDR and the Expr of its corresponding ZGA gene (2-cell vs zygote, fold change ≥ 4, FDR < 0.01; Supplementary Table 2) across single cells during the transition from the zygote to the 2-cell stage. We found that 338 promoter NDRs and 7822 distal NDRs were positively linked to 301 and 2239 ZGA genes, respectively, while 356 promoter NDRs and 2728 distal NDRs were negatively linked to 317 and 1226 ZGA genes, respectively (Fig. 4a-b; Supplementary Table 3). Although the Met in those CREs was not significantly associated with ZGA gene expression (data not shown), the overall Met levels of these positively correlated CREs were lower in 2-cell embryos than in zygotes (Extended Data Fig. 5a and c). Notably, the Acc of these positively correlated CREs was specifically increased in each individual cell in 2-cell embryos, but this increase was accompanied by a drastic global Acc decrease during this period (Extended Data Fig. 5b and d). These results suggest that robust chromatin reprogramming occurs during ZGA to remove regulatory memory from gametes and rebuild the zygotic regulatory network.

**Figure 4.**
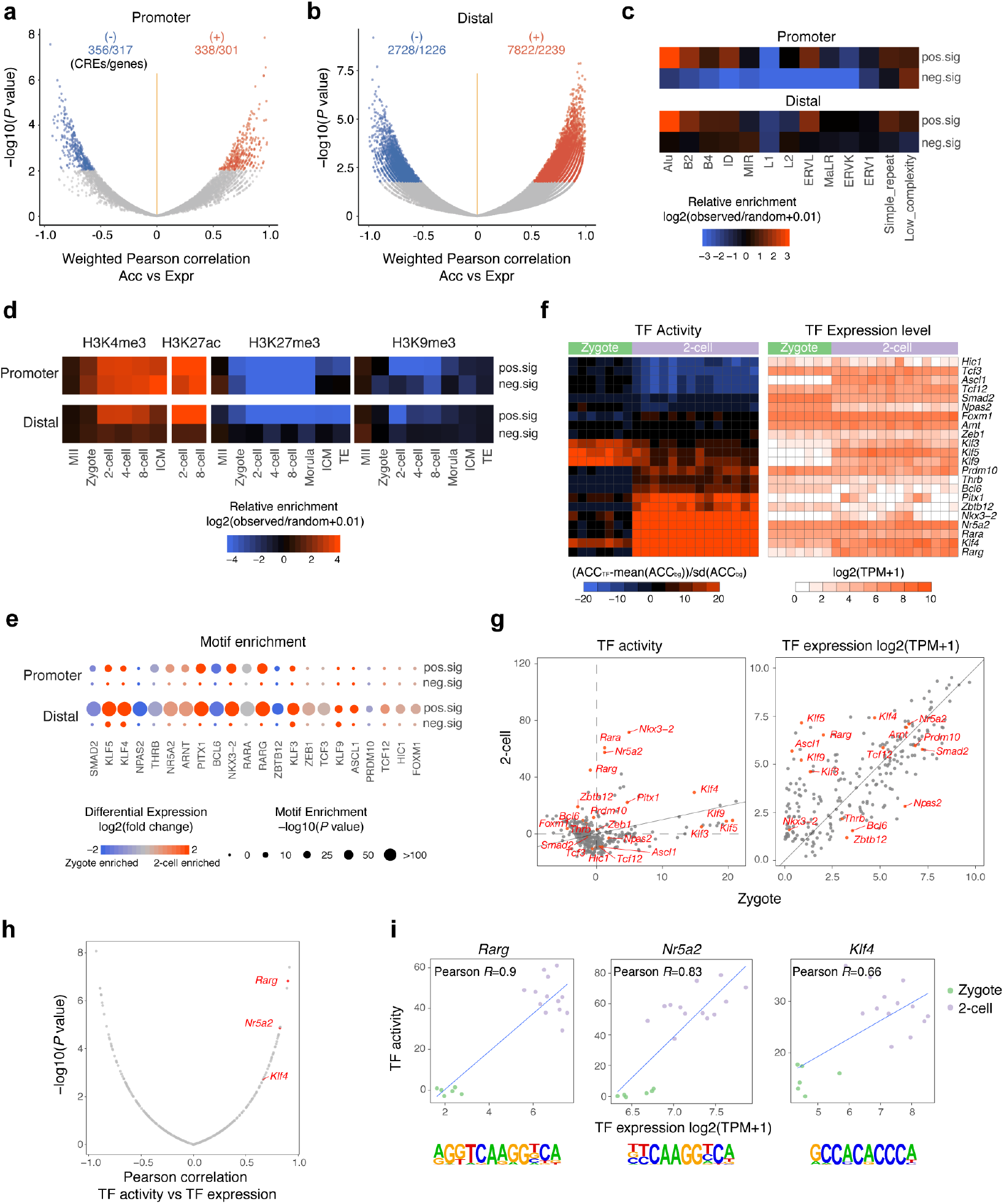
scNOMeRe-seq reveals a ZGA associated regulome. (**a-b**) Volcano plot showing the weighted Pearson correlations between chromatin accessibility of promoter-NDRs (**a**) / distal-NDRs (**b**) and the expression level of corresponding ZGA genes across cells from zygote to 2-cell stage. Significant associations (FDR < 0.1) are in red (positive) and blue (negative). The number of CREs (CRE, the NDR that was significantly correlated with ZGA gene) and unduplicated genes are labeled. (**c**) Heat map showing the enrichment of ZGA associated CREs in repeats. The enrichment was calculated as log2 ratio for the numbers of observed CREs that overlap with repeats divided by the numbers of random regions that overlap with repeats. (**d**) Heat map showing the enrichment of ZGA associated CREs in histone modifications of early embryos from available published data (ref. 7-9). The enrichment was calculated as log2 ratio for the number of observed CREs that overlap with histone modification peaks divided by the number of random regions that overlap with histone modification peaks. (**e**) TF motifs identified from ZGA associated CREs. Only TFs with the *P* value < 1 x 10^-10^ and TPM ≥ 5 at least at one stage were included. The color of point represents the differential gene expression level between 2-cell stage and zygote. The size of point represents the significance of motif enrichment. (**f**) Heat map showing the TF activity (left) and expression level (right) of ZGA enriched TFs in each individual cell of zygote and 2-cell embryos. (**g**) Scatter plot showing the difference of TF activity (left) and expression level (right) between zygote and 2-cell stage. The genes labeled in red indicate the TFs showing significantly differential activity (left, FDR < 0.1) or expression level (right, FDR < 0.01). (**h**) Volcano plot showing the Pearson correlations between TF activity and expression level across cells from zygote to 2-cell stage. Significantly correlated (FDR < 0.1) TFs that enriched in ZGA associated CREs and showed higher TF activities and expression level in 2-cell embryos are labeled. (**i**) Scatter plot showing TF activity and expression level of *Rarg*, *Nr5a2* and *Klf4* in each single cell of zygote and 2-cell embryos. The corresponding enriched DNA-binding motifs are shown at the bottom of each TF.

To explore how ZGA is regulated in early embryos, we further comprehensively analyzed the enrichment of repeat elements and histone modifications in ZGA-associated CREs. The positively correlated CREs, but not the negatively correlated CREs, were preferentially enriched with Alu, B2, B4 and ERVL repeat classes as well as active histone modifications (H3K4me3 and H3K27ac) in both promoter and distal regions (Fig. 4c-d). H3K4me3 was gradually established at the majority of positively correlated CRE loci from the MII oocyte stage to the 2-cell stage and was colocalized with H3K27ac in both promoter and distal regions, while repressive histone modifications (H3K27me3 and H3K9me3) were gradually removed from these regions (Extended Data Fig. 6a-c). Moreover, the positively correlated CREs were clustered in regions enriched with active histone modifications and deficient in repressive histone modifications, implying a high-dimensional regulatory structure of ZGA CREs (Extended Data Fig. 6d-e). Furthermore, we investigated which TFs might be responsible for the establishment of ZGA-associated CREs. Notably, both positively correlated promoter CREs and distal CREs were highly enriched with *Arnt*, *Bcl6*, *Klf5*, *Nkx3-2*, *Nr5a2*, *Rara*, *Rarg*, *Pitx1*, and *Thrb* motifs; however, the negatively correlated CREs showed no enrichment with TFs in either promoter or distal regions (Fig. 4e). Considering the global decreases in Acc during the ZGA process and the characteristics of the negatively correlated CREs described above, these negative correlations seemed to simply reflect global changes in Acc rather than representing repressive regulation during ZGA. We next calculated TF activity (see Methods) in each individual cell (Fig. 4f). Notably, we found that *Klf4*, *Nkx3-2*, *Nr5a2* and *Rarg* showed high TF activity and high expression levels in 2-cell embryos compared to zygotes (Fig. 4g). More importantly, the TF activity of *Rarg*, *Nr5a2* and *Klf4* was strongly positively correlated with the expression levels of these genes, further supporting their potential roles in regulating ZGA-associated CREs (Fig. 4h-i). It is worth noting that among these three TFs, *Klf4* already showed high expression levels and high TF activity at the zygote stage, while both *Rarg* and *Nr5a2* showed almost no TF activity at the zygote stage, implying that *Klf4*, as a maternal factor, might contribute to initiating the ZGA process as early as the zygote stage (Fig. 4f and i).

### Mutually exclusive regulome confers ICM/TE lineage segregation

Along with gradual increases in heterogeneity among blastomeres in preimplantation embryos, establishment of cell lineage-specific transcription regulatory networks occurred beginning in unified totipotent zygotes that generated ICM and TE cells to enable further embryo development. To reveal the potential active CREs during this process, we determined the correlations between the Acc of each promoter/distal NDR and the Expr of its corresponding ICM/TE-specific expressed genes (specifically expressed in ICM: 766 genes, TE: 930 genes; Supplementary Table 4) across single cells during preimplantation development (Fig. 5a-b). The NDRs significantly correlated with ICM-or TE-specific expressed genes were termed ICM.CREs (positive: 497 in promoters, 4086 in distal regions; negative: 210 in promoters, 1559 in distal regions) or TE.CREs (positive: 774 in promoters, 5109 in distal regions; negative: 424 in promoters, 3445 in distal regions), respectively (Fig. 5a-b; Supplementary Table 5). Consistent with the ZGA-associated CREs, the positively correlated ICM/TE CREs also showed strong enrichment for active histone markers and depletion of repressive histone markers (Extended Data Fig. 7a-b). Notably, all of the known enhancers for three key ICM/TE TFs (*Pou5f1*, *Nanog*, and *Cdx2*) that we analyzed were revealed to be present in preimplantation embryos or in embryonic stem cells, confirming that the CREs identified by our correlation analysis could cover known active enhancers (Fig. 5c-d, Extended Data Fig. 7c-f)^32–34^. Specifically, we found three positively correlated CREs (#2, #3, and #4) corresponding to three known enhancers of *Pou5f1*; importantly, CRE #4 showed the highest positive correlation coefficient in our analysis, consistent with previous findings that the known enhancer corresponding to CRE #4 is the dominant enhancer regulating *Pou5f1* expression during preimplantation development (Fig. 5c-d)^32^. Thus, these results validate the accuracy of our analysis.

**Figure 5.**
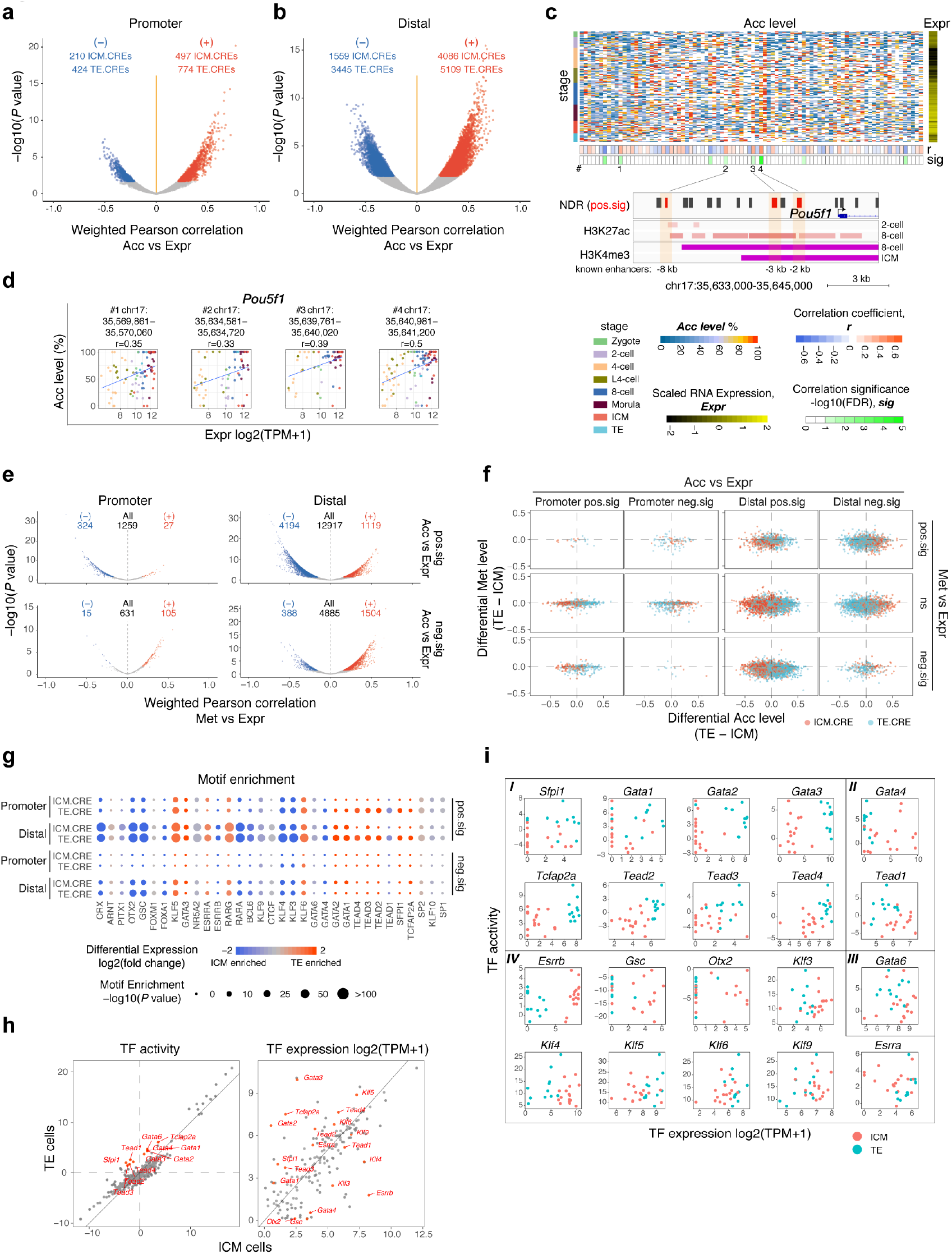
Mutually exclusive regulome confers ICM/TE lineage segregation. (**a-b**) Volcano plot showing the weighted Pearson correlations between chromatin accessibility of NDRs at promoter (**a**) / distal (**b**) regions and the expression level of corresponding ICM/TE specific expressed genes across preimplantation development. Significant associations (FDR < 0.1) are in red (positive) and blue (negative). ICM.CRE, the NDR that was significantly correlated with ICM-specific-expressed gene; TE.CRE, the NDR that was significantly correlated with TE-specific-expressed gene. The number of CREs are labeled. (**c**) (top) Heat map showing chromatin accessibility (Acc level) of *Pou5f1* gene locus surrounding NDRs (from TSS upstream 100 kb to TES downstream 100 kb), and expression level (Expr, scaled log2(TPM+1)) of *Pou5f1* in each individual cell of early embryos; the weighted Pearson correlation coefficients (r) and significance (sig) are shown. #, the labels of positive-correlated CREs. (bottom) IGV snapshot showing the distribution of *Pou5f1* surrounding NDRs (positive-correlated CREs are in red), H3K27ac and H3K4me3 peaks. Three known enhancers of *Pou5f1* are shaded. (**d**) Scatter plot showing the expression level of *Pou5f1* and chromatin accessibility of positive-correlated CREs which have been labeled in (**c**). The genomic coordinates of CREs and the correlation coefficients are shown. (**e**) Volcano plot showing the weighted Pearson correlation between DNA methylation level and gene expression level of positive-correlated (top) and negative-correlated (bottom) CREs. (**f**) Scatter plot showing the differential chromatin accessibility and DNA methylation level of ICM.CREs and TE.CREs between ICM and TE cells at promoter (left) and distal (right) regions. Positive value indicates a higher epigenetic level in TE cells, negative value indicates a higher epigenetic level in ICM cells. (**g**) TF motifs identified from ICM.CREs and TE.CREs. Only TFs with the *P* value < 1 x 10^-10^ and TPM ≥ 5 at least at one stage were included. The color of points represents the differential gene expression level between ICM and TE. The size of point represents the significance of motif enrichment. (**h**) Scatter plot showing the difference of TF activity (left) and expression level (right) between ICM and TE. The genes labeled in red indicate the TFs showing significantly differential activity (left, FDR < 0.1) or expression level (right, FDR < 0.01) between ICM and TE. (**i**) Scatter plot showing the TF activity and expression level in each individual cell of ICM and TE. Ι, the TFs showing higher activity and expression level in TE cells; Ⅱ, the TFs showing higher activity in TE cells but showing higher expression level in ICM cells; Ⅲ, the TF showing higher activity in TE cells but showing no difference in gene expression level; Ⅳ, the TFs showing no difference in activity but showing higher expression level in ICM cells.

To explore how ICM/TE-associated regulatory networks are regulated in early embryos, we comprehensively analyzed different epigenetic molecular layers in these CREs. First, we calculated the correlations between Met and Expr for each CRE-gene pair. Interestingly, we observed mainly negatively correlated Met vs Expr CRE-gene pairs (27 positive vs 324 negative pairs in promoters; 1119 positive vs 4194 negative pairs in distal regions) for the positively correlated Acc/Expr CREs, while we observed mainly positively correlated Met vs Expr CRE-gene pairs (105 positive vs 15 negative pairs in promoters; 1504 positive vs 388 negative pairs in distal regions) for the negatively correlated Acc/Expr CREs (Fig. 5e). These results not only reveal the complex interplay between Met and Acc in regulating ICM/TE lineage-associated regulatory networks during preimplantation development but also confirm our speculation that the CREs with Acc/Expr positive and negative correlations might be bound by activators and repressors, respectively. Regardless, the Met levels of both ICM.CREs and TE.CREs were lower in TE cells than in ICM cells, reflecting the more extensive erasure of genome-wide Met in TE cells (Fig. 5f). Next, we measured the dynamics of different molecular layers in these CREs during preimplantation development. Clearly, Acc and active histone modifications (H3K4me3 and H3K27ac) gradually increased in both positively and negatively correlated Acc/Expr CREs beginning at the 2-cell stage, while repressive epigenetic modifications (Met, H3K9me3 and H3K27me3) were already depleted in zygotes and remained depleted throughout preimplantation development in these regions, suggesting an overall priming of the epigenetic environment during ICM/TE lineage differentiation (Extended Data Fig. 8a-f). Interestingly, we also noticed that ICM.CREs were activated earlier than TE.CREs, as we observed clearly higher levels of active epigenetic modifications in ICM.CREs than in TE.CREs at the 2-cell stage (Extended Data Fig. 8a, b, e and 9a-b). Subsequently, TE.CREs were quickly activated and showed higher levels of active epigenetic modifications than ICM.CREs at the 8-cell stage, suggesting a stepwise activation of ICM.CREs and TE.CREs during preimplantation development (Extended Data Fig. 8a, b, e and 9a-b).

Finally, we investigated which TFs might be responsible for the establishment of differential regulatory networks in ICM and TE lineages. Notably, we found that the commonly enriched TFs in positively correlated ICM/TE CREs were ZGA drivers that showed high TF activity as early as the 2-cell embryo stage, such as *Nr5a2*, *Rarg*, *Rara*, *Bcl6*, etc., indicating that the earliest initiation of both ICM and TE programs occurs during the ZGA process (Fig. 5g and Extended Data Fig. 9c). In addition, three TFs, *Crx*, *Arnt* and *Pitx1,* were more enriched in ICM.CREs, while *Ctcf*, *Klf3/4/6/9/10*, *Gata1/2/4/6*, *Tead1/2/3/4*, *Sfpi1*, *Tcfap2a* and *Sp1/2* were more enriched in TE.CREs (Fig. 5g). Interestingly, we observed that TE lineage-specific TFs, such as *Tcfap2a* and *Gata* family TFs, were enriched in the negatively correlated ICM.CREs, suggesting their repressive roles in regulating the ICM program (Fig. 5g). Moreover, most TE.CRE-specifically enriched TFs showed higher activity and expression levels in TE cells than in ICM cells, while ICM.CRE-associated TFs showed higher expression levels in ICM cells than in TE cells (Fig. 5h-i and Extended Data Fig. 9c). Together, these results suggest that a mutually exclusive regulatory network is adopted to gradually establish and stabilize the different ICM and TE lineage fates, especially for the TE lineage; specific drivers of this lineage establish a TE program while repressing the ICM program, forcing the TE lineage to separate from the ICM lineage.

## Discussion

In conclusion, we have developed a single-cell multiomics sequencing technology, scNOMeRe-seq, that can be used to profile transcriptomes, DNA methylomes and Acc in parallel in the same individual cell with high accuracy, sensitivity and genome coverage. Taking advantage of this powerful tool, we have also characterized multiple molecular layers of mouse preimplantation embryos at single-cell resolution and have explored the associations between different epigenome layers and transcriptional output, providing new insights to enhance functional understanding of epigenetic reprogramming during mouse preimplantation development. Specifically, our results reveal that PG blastomeres show delayed development and abnormal DNA methylomes, in contrast to aneuploid blastomeres. The changes in Acc not only reflect the dynamic regulatory landscape but also may be substantially derived from asynchronous cell cycles of blastomeres that are irrelevant with transcriptional regulation, highlighting the importance of functional interpretations of epigenetic reprogramming at single-cell resolution. Using the DNA methylomes of all the individual cells within individual embryos, we reconstructed genetic lineages from zygotes to 8-cell embryos and revealed that asymmetric cleavage might be the major driver of the gradual increases in transcriptome heterogeneity among blastomeres that occur during the first three cleavages. Despite global demethylation in early embryos, allele-specific Met patterns inherited from oocytes and sperm are maintained throughout preimplantation development. The associations between Acc/Met and Expr at promoter regions in single cells are consistent with the findings from bulk samples; however, the positive correlations between Met and Expr at gene body regions are largely inherited from maternal genomes and are absent in the paternal genomes of early embryos. The Acc of parental memory-related regions appears to be substantially erased during the ZGA process and reconfigured in concert with the influences of histone modifications, Met, repeats, TFs, and possible high-dimensional chromatin structures to ensure proper activation of the zygotic genome. The overall-primed ICM/TE lineage-associated CREs are partially activated as early as the 2-cell embryo stage and are asynchronously activated in the following preimplantation stages. Intriguingly, TE lineage-specific TFs seem to play dual roles in activating the TE program and repressing the ICM program, thereby segregating the TE fate from the ICM fate. Taken together, our findings not only provide new insights into the functional regulatory landscape in preimplantation development but also elucidate the fundamental mechanisms of epigenetic regulation.

## Supporting information

Supplemental sections

## Methods

Methods and any associated references are available in the supplemental sections.

## Acknowledgements

This work was supported by the grants from National Key Research and Development Program (2018YFC1004000, 2018YFC1004101 and 2017YFA0105001), National Natural Science Foundation of China (31571544, 81730038, 81521002 and 31701261) and the Fundamental Research Funds for the Central Universities-Peking University Clinical Scientist Program. Y.W. was supported by Postdoctoral Fellowship of Peking-Tsinghua Center for Life Science and the grant from China Postdoctoral Science Foundation (2016M600873).

## Author contributions

Y.W. conceived the method. L.Y. and J.Q. conceived the project. Y.W. and P.Y. performed experiments with the help of M.Y., Y.H., Y.N., X.Z. Y.W., Z.Y., and Y.P. performed computational analysis. Y.W., P.Y. and Z.Y. wrote the manuscript with feedback from all authors.

