## Supplemental sections for "Single-cell multiomics sequencing reveals the functional regulatory landscape of early embryos"

### Supplementary Information

#### Embryo collection

All animal-related experimental procedures were carried out under ethical guidelines set forth by the Animal Care and Use Committee of Peking University Health Science Center. The female mice used in this study were 6- to 8-week-old B6D2F1/J (BDF1) mice, and the male mice were 12-week-old 129S1 mice. All mice were in good health. For mouse preimplantation embryo collection, female mice were superovulated by injection of 7.5 IU of pregnant mare serum gonadotropin (PMSG) followed by 7.5 IU human chorionic gonadotropin (hCG) 42-48 h later. Immediately after hCG injection, the female mice were placed in a cage with males. After vaginal plugs appeared, zygotes were collected from the mouse oviducts, and the embryos were further cultured in KSOM medium (Zenith Biotech). Embryos at different stages were collected 24-26 h (zygotes), 36-40 h (2-cell embryos), 52-54 h (4-cell embryos), 58-60 h (late 4-cell embryos), 66-70 h (8-cell embryos), 78-82 h (morulae) and 94-98 h (blastocysts) after hCG injection. The embryos were exposed to acidic solution (0.5% concentrated HCl) to remove the zona pellucidae and were then washed with 0.5% HSA/DPBS (Vitrolife, Gibco) to remove polar bodies and somatic cells. The embryos were further dissociated into single cells by incubating them in a trypsin (Sigma-Aldrich) and Accutase (Millipore) solution (at a 1:1 volume ratio) for 15-50 min. Then, the embryos were washed with 0.5% HSA/DPBS three times.

#### Lineage tracing with FITC injection

One blastomere of each 2-cell stage mouse embryo was microinjected with FITC-coupled dextran (Sigma-Aldrich) to indicate its daughter cells at the 4-cell stage or its granddaughter cells at the 8-cell stage. To indicate which blastomeres of 8-cell embryos were from the same mother cells of 4-cell embryos, one blastomere of each 4-cell stage embryo was microinjected with FITC-coupled dextran. After those embryos developed to the desired stage, each embryo was dissociated into single cells as mentioned above. The FITC-labeled cells were identified under a fluorescence microscope before library preparation. All available single cells were processed for scBS-seq[35].

**Library preparation and sequencing**

Single cells were picked by mouth pipetting, and each was transferred into a 200 μl PCR tube containing 3 μl of cell lysis buffer (1x RT buffer (Invitrogen), 0.5% NP40, 5 mM DTT, 2 U RNase OUT (Invitrogen), and 0.2 μl of magnetic beads (Invitrogen, cat. # 65001)). The cells were lysed on ice for 10 min and vortexed for 30 sec. Then, the tubes were placed on a magnet for 5 min to separate the beads (containing the intact nuclei) and supernatant (containing the RNA transcripts). The supernatant was transferred to a new PCR tube, and RNA libraries were prepared following the MATQ-seq protocol[28]. External RNA Controls Consortium (ERCC) spike-ins (Ambion) were added to the supernatant at dilutions of 1:10000-100000. For the split RNA libraries, the supernatant was split into 2 aliquots before first-strand synthesis, and the libraries were prepared separately. In parallel, the precipitated beads (containing the intact nuclei) were resuspended in 5 μl of GpC methylase reaction buffer (1x M.CviPI reaction buffer (NEB), 5 U M.CviPI (NEB), 160 μM S-adenosylmethionine (NEB), 0.25 mM EDTA (Thermo), 0.25 mM phenylmethylsulfonyl fluoride (Sigma-Aldrich), and 1 pg of lambda DNA (NEB)) and incubated in a thermocycler at 37 °C for 60 min to methylate GpC before heat inactivation for 25 min at 65 °C. After *in vitro* GpC methylation, 0.5 μl of protease (Qiagen, 20 mg/ml) and 10 ng of carrier RNA (Qiagen) were added into the mixture, which was incubated at 50 °C for 3 h in a thermocycler to release genomic DNA before heat inactivation at 75 °C for 30 min. Then, the genomic DNA was bisulfite-converted (Thermo), and the scBS-seq protocol was followed to prepare DNA libraries[35]. For the FITC-tracing experiments, the dissociated single cells were transferred into a 200 μl PCR tube containing 5 μl of RLT plus (Qiagen) to release genomic DNA, and then the scBS-seq protocol was followed to prepare DNA libraries. All the libraries in this study were sequenced on an Illumina HiSeq X Ten platform with 150 bp paired-end reads. With scNOMeRe-seq, we detected 3.49 million unique WCG sites, 31.04 million unique GCH sites and the expression of 14416 (TPM>0) GENCODE genes per cell on average.

**scRNA-seq data processing**

The raw reads were trimmed with Trim Galore (v0.4.4) to remove the primer sequences and low-quality bases (parameters: trim_galore --paired --quality 20 --phred33 --stringency 3 --gzip --length

36). The trimmed reads were aligned to the GENCODE NCBIM37 reference genome (corresponding to the University of California, Santa Cruz (UCSC) mm9 genome) with STAR software[36] with the default settings, and only the unique mapped reads with MAPQ values > 50 were retained for further analysis. The reads mapped to rRNA were filtered out with RSeQC[37]. The coverages of the transcripts were evaluated with RSeQC. The reads mapped to each gene were counted with featureCounts[38] (parameters: featureCounts -p -t exon -g gene_id), and the gene expression levels were calculated using TPM values. Samples were discarded for subsequent Expr analysis that had (1) less than 9500 genes detected, (2) the library size less than 0.1 million counts, (3) reads for mitochondrial genes accounted for over 30%, or (4) reads with a fraction of the top 50 features over 40%. Then, the split cell libraries that passed quality control were merged using the *DESeq2*[39] R package. In total, 221 single-cell RNA libraries were retained in this study.

**Principal component analysis and hierarchical clustering of individual cells using RNA expression data**

PCA and hierarchical clustering were performed to analyze cell populations with the Expr data. Genes expressed in fewer than 6 cells were discarded. Pairwise Spearman correlation coefficients were calculated. PCA was performed using the *pcaMethods*[40] R package. The *hclust* function with the *ward.D2* method was used for unsupervised hierarchical clustering.

**SC3 clustering**

To assign single cells from the blastocyst stage into ICMs and TEs and to identify markers of the two types of cells, we used the *SC3*[41] R package on the scRNA-seq data. The quality-controlled expression matrix was passed into *SC3,* and the clusters were plotted with the sc3_plot_consensus function. The marker genes were further identified with the sc3_plot_markers function.

**Preimplantation embryo developmental pseudotime analysis**

Although PCA clustered the individual cells into their biological developmental stages, the cells within each group were not ordered by their developmental time points. Therefore, individual cells were further ordered with the *destiny*[42] R package using the top 2000 variable genes in the individual cells.

**Differential RNA expression**

Differential Expr between groups and stages was analyzed using the R package *DESeq2*.

**Copy number variation estimation using scRNA-seq data**

To evaluate the CNV effect on Expr, we inferred the CNV status of each cell by averaging the expression of the ordered genes across the genome, as previously described[43]. In brief, genes were ordered along the genome by their genomic locations, and the expression of each gene was adjusted to the average expression of the 50 upstream and downstream genes. The adjusted expression values were further centered to infer the CNV status.

**scNOMe-seq and scBS-seq data processing**

The single-cell Met data was analyzed as previously described[44]. The paired-end FASTQ reads were processed as two single-end FASTQ files because chimeric reads were produced during library preparation. The raw paired-end FASTQ reads were trimmed to remove the first 11 bp of the random primer sequences, Illumina adapter sequences and low-quality bases with Trim Galore (v0.4.4) in single-end mode (parameters: trim_galore --clip_R1 11 --quality 20 --stringency 3 --length 30). The trimmed reads were aligned to the lambda and UCSC mm9 genome with Bismark[45] in single-end mode (parameters: bismark --bowtie2 --non_directional). The single-end-mode mapped reads were merged, and PCR duplicates were removed (PICARD) for downstream analysis. The bismark_methylation_extractor function was used to call the methylation value for each cytosine site in the genome (parameters: bismark_methylation_extractor -s --multicore 4 --gzip --cytosine_report --CX), which required at least 1x coverage at the cytosine sites. WCG (W includes A and T) and GCH (H includes A, C, and T) Met levels were calculated to represent the Met levels and Acc levels, respectively. The samples with less than 0.5 million covered WCG and 5 million covered GCH sites were discarded before downstream analysis. In total, 218 scNOMe-seq and 26 scBS-seq libraries passed the quality control steps.

**Quantification of DNA methylation and accessibility**

The Met and Acc levels were calculated as the sum of methylated reads (C) divided by the total covered reads (sum of the methylated reads and unmethylated reads (T)) for each WCG and GCH site, respectively. The Met and Acc levels of the genomic regions and single cells were measured as the average WCG and GCH levels, respectively. The TSS Met level was measured as the average WCG level within 1 kb upstream and 0.5 kb downstream of the TSS. TSS Acc was measured as the average GCH level within 200 bp upstream and 100 bp downstream of the TSS.

**Nucleosome-depleted region identification**

NDRs were defined as previously described[4]. Briefly, the GCH data of each single cell from the same developmental stage were first aggregated. The number of C and T reads in the GCH context within a 120 bp window with 20 bp spacing was calculated, and the significance of the difference from the genomic background was analyzed with the Chi-square test. Regions with significantly elevated GCH methylation with $P$-values $\leq 10^{-15}$, lengths $\geq$ 140 bp and covered GCH sites $\geq$ 5 were defined as NDRs. The NDRs that overlapped with the ENCODE blacklist (mm9, http://mitra.stanford.edu/kundaje/akundaje/release/blacklists/) were removed before downstream analysis. The defined NDRs were further classified as Promoter_NDRs (within 2 kb of the TSS; NDRs that overlapped with the TSS termed TSS_NDRs, while those that did not overlap with the TSS were termed Proximal_NDRs) and Distal_NDRs (at least 2 kb away from the TSS) in downstream analysis.

**Copy number variation detection using scBS-seq data**

To detect the CNV in each cell based on the scBS-seq data, we inferred the CNVs using the HMMcopy[46] R package. First, the sequenced reads were counted with readCounter in 1 Mb bins across the mouse genome, and the read count of each bin was adjusted by correcting for the GC content and genomic mappability. Then, the median adjusted read counts of the consecutive bins were used to infer the CNVs of the genomic regions. Median values of more than 1.36 and less than 0.67 observed in >10 consecutive bins were defined as gains and losses, respectively.

**Principal component analysis and hierarchical clustering of blastomere DNA methylation and chromatin accessibility**

To analyze cell populations with the Met and Acc datasets, we performed PCA and hierarchical clustering using the WCG levels of genome-wide 5 kb tiles and the GCH levels of merged NDRs. Genomic regions that covered fewer than 3 sites were discarded. Spearman correlation coefficients were calculated with the parameter *pairwise.complete.obs*. PCA was performed using the *pcaMethods* R package. The *hclust* function with the *ward.D2* method was used for unsupervised hierarchical clustering.

**Profiling of DNA methylation and chromatin accessibility**

To profile the Met and Acc levels around genic regions (from 2.5 kb upstream of the TSS through the gene body to 2.5 kb downstream of the transcription end site (TES)), the average GCH levels and WCG levels within predefined running windows were computed for each gene in each single cell, as previously described[23]. The running window was defined as a 150 bp window with a 50 bp step for the TSS upstream and the TES downstream. The gene body (from the TSS to TES) of each gene was divided into 100 equal fractions, and the running window was defined as a 2-fraction window with a 1-fraction step. To profile the Met and Acc levels around NDRs (from 3 kb upstream of each NDR to 3 kb downstream), the average GCH levels and WCG levels within a 150 bp running window with a 50 bp step were computed for each NDR in each single cell. The values of all the same genomic locations from individual cells were combined to plot the average profile for single cells. The values from cells at the same stage or in the same group were combined to plot the average profile for the stage or group, respectively.

**Correlation analysis**

To profile the relationship between Met/Acc and Expr around the genic regions, the Pearson correlation was calculated between the average WCG/GCH level within the running window (described above) and the Expr of its corresponding gene across different genes in a single cell. To calculate the associations between Met/Acc and Expr at specific genomic regions (the TSS and gene body), the Pearson correlation was calculated between the average WCG/GCH level within the region and the Expr across different genes in a single cell.

To infer the functional CREs with associated genes, we computed the correlation between the dynamic Met/Acc level of each NDR and the expression of its corresponding gene across cells during preimplantation development. All possible relationships between NDRs and genes within 100 kb of the gene (upstream of the TSS and downstream of the TES) were considered. NDRs with a coverage of fewer than 3 sites in a single cell were filtered out. NDRs covered in less than 25% of the cells and nonvariable NDRs were discarded. Genes expressed at low levels (expressed (TPM>0) in fewer than 5 cells) and nonvariable genes were discarded. We calculated a weighted Pearson correlation coefficient (using the unique WCG or GCH sites covered within the NDRs in each single cell as a weight) and tested the significance of the coefficients with two-tailed Student's t-tests. The *P*-values were further adjusted by the Benjamini-Hochberg approach.

**Allele-specific analysis of RNA expression, DNA methylation and accessibility**

We downloaded the SNPs of 129S1, C57BL6NJ and DBA2J from the website of the Mouse Genome Project (ftp://ftp-mouse.sanger.ac.uk/REL-1211-SNPs_Indels/). Only the SNPs that could distinguish the paternal (129S1) and maternal (C57BL6NJ X DBA2J) genomes were used in our analysis. For allele-specific analysis of the RNA-seq data, 172319 SNPs within exonic regions were used to split the mapped reads. For the scBS-seq data, 896161 SNPs (excluding SNP sites with C or T) in the whole genome were used to split the mapped reads into paternal- and maternal-origin reads. The split reads were further processed to calculate the paternal and maternal allele-specific gene expression, Met and Acc levels. To profile the allelic Met and Acc levels around genic regions (from 2.5 kb upstream of the TSS through the gene body to 2.5 kb downstream of the TES), the average allelic WCG levels and GCH levels within predefined running windows (500 bp window with a 100 bp step for the TSS upstream and the TES downstream; 10-fraction window with a 2-fraction step for the gene body (100 equal fractions)) were computed for each gene in each single cell. Local correlations were calculated between the average allelic WCG/GCH level within the running window and the total Expr of its corresponding gene across different genes in a single cell.

The epigenetic modification levels of 500 bp tiles were extracted from the parental alleles of each single cell separately. The tiles retained in each stage were required to cover at least 3 cells per allele. The numbers of methylated sites (C) and unmethylated sites (T) were summed in each 500 bp tile per allele for each stage. Fisher's exact test was used to examine the differences between two alleles for each tile. Tiles with FDR values less than 0.01 were considered to differ significantly between alleles.

**Integrative analysis of public data and resources**

Repetitive elements (such as LINEs, SINEs, Alu elements, etc.) and related genomic annotations (such as the TSS, TES, gene body, etc.) were downloaded from the UCSC. Promoters were defined as the regions from -1.5 kb to +0.5 kb relative to the TSS. Histone modification data for mouse preimplantation embryos were downloaded from the Gene Expression Omnibus (GEO) database (GSE71434 for H3K4me3 modifications; GSE97778 for H3K9me3 modifications; GSE72784 for H3K27ac modifications; GSE73952 for H3K27me3 modifications) and integrated into our analysis. The peaks were downloaded from the GEO database. The signal intensity of each histone modification was calculated with an identical procedure. In detail, the downloaded raw FASTQ reads were trimmed with Trim Galore (parameters: trim_galore --paired --quality 20 --phred33 --stringency 3 --length 36). The processed clean reads were mapped to the UCSC mm9 genome using BWA with the default settings[47]. The duplicated reads were removed with PICARD software, and only unique mapped reads with MAPQ values ≥ 30 were retained. The sequencing coverage was further normalized, and the histone modification signal intensity of the investigated regions was calculated with the *Deeptools* package[48].

**Transcription factor analysis**

To find the enriched TF motifs in different CRE data sets, findMotifsGenome.pl in HOMER[49] was used (parameters: -size given -len 8,10,12). Only TFs with the $P$ value $< 1 \times 10^{-10}$ and TPM ≥ 5 at least at one stage were included.

To further infer the TF activity in each single cell, we calculated background-corrected z-scores for the average Acc of the TF-binding sites (TFBSs) for each TF in each single cell. A similar strategy has been developed for single-cell ATAC-seq[50]. More precisely, all stage-defined distal NDRs were scanned to find TFBSs using FIMO[51]. Position frequency matrices were converted from 413 Homer motifs. We kept the motif TFBSs with *P*-values less than $1 \times 10^{-4}$. The overlapping motifs for each TF were merged. The average Acc of the TFBSs (extended from the center to +- 50 bp) in each single cell for each TF was calculated as the raw TF activity ($ACC_{TF}$). Then, we permuted the data 1000 times for the TFBSs for each TF in each single cell and calculated the average Acc ($ACC_{bg}$) for each permutation. For each TF in each cell, TF activity was calculated as ($ACC_{TF}$-mean($ACC_{bg}$))/sd($ACC_{bg}$).

**Data availability**

All sequencing data in the current study have been deposited in the GEO with the accession number GSEXXXX.

**Code availability**

All the computational code used in this study for data processing and analysis are available upon requesting from corresponding authors.

**a** **b** **c** **d** **e** **f**

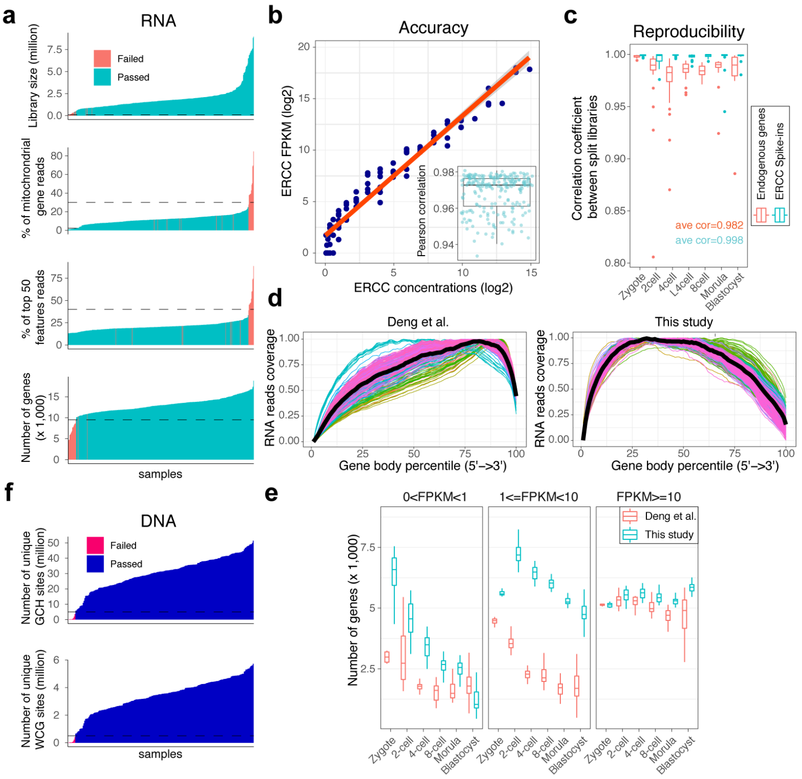


**g**

| | Zygote | 2-cell | 4-cell | L4-cell | 8-cell | Morula | Blastocyst | total |
|---|---|---|---|---|---|---|---|---|
| RNA passed/ DNA failed | 1 | 5 | 2 | 2 | 1 | 0 | 0 | 11 |
| RNA failed/ DNA passed | 0 | 0 | 0 | 0 | 0 | 0 | 8 | 8 |
| Both failed | 0 | 0 | 1 | 0 | 0 | 0 | 3 | 4 |
| Both passed | 12 | 19 | 45 | 22 | 31 | 24 | 57 | 210 |

**h** **i**

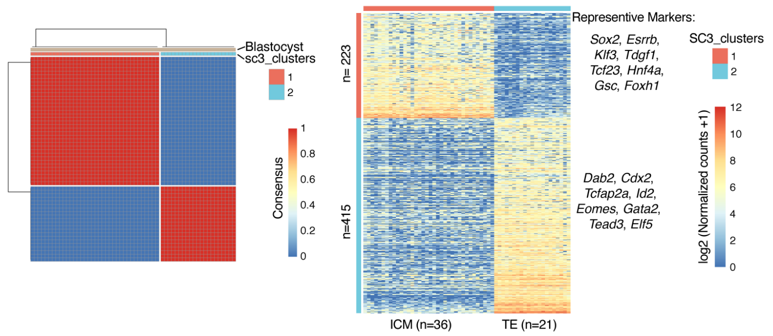


**Extended Data Figure 1 Quality control of the scNOMeRe-seq datasets.**

(**a**) Quality of RNA libraries. Cells with below the threshold of library size (total number of counts, cutoff 0.1 million), number of expressed genes (log2 normalized read counts > 0, cutoff 9,500) were discarded. Cells with high fraction of mitochondrial gene reads (cutoff 30%), fraction of top 50 features reads (cutoff 40%) were discarded.

(**b**) Scatter plot showing the relationship between expected ERCC spike-ins concentrations and detected expression level (FPKM, log2) of a random selected single cell, as an example. Box plot at the right bottom showing the Pearson correlation coefficients between expected ERCC spike-ins concentrations and detected expression level of all cells passed quality control.

(**c**) Box plot showing the Pearson correlation coefficients of expression level (FPKM, log2) of endogenous genes or ERCC spike-ins between two split libraries of a single cell.
(**d**) Profiles of RNA reads coverage along gene bodies in previously published data (left, ref. 29) and this study (right). Black thick lines showing the average rates of all cells.
(**e**) Box plot showing the gene numbers at low, medium and high expression level of each single cell at each stage in previously published data (ref. 29) and this study.
(**f**) Quality of DNA libraries. Cells with below the threshold of unique GCH sites (cutoff 5 million) or unique WCG sites (cutoff 0.5 million) were discarded.
(**g**) Summary of cell numbers of quality control results in this study. Split RNA libraries from the same cell were merged, counting as one RNA library.
(**h**) Consensus plot of E3.5 blastocyst with gene expression level (log2 normalized counts). Cells are clustered into two stable clusters.
(**i**) Heat map showing gene expression level of ICM, TE lineage specific markers for cluster 1 and 2, respectively. Representative markers are shown, with $P$ value < 0.01 and AUROC > 0.85.

**a**

| | Zygote | 2-cell | 4-cell | L4-cell | 8-cell | Morula | ICM | TE | total |
|---|---|---|---|---|---|---|---|---|---|
| Normal | 8/1 | 12/4 | 37/2 | 19/1 | 22 | 16 | 20 | 15 | 149/8 |
| PG | 4 | 3/1 | 4 | 3/1 | 7/1 | 0 | 0 | 0 | 21/3 |
| Aneu | 0 | 4 | 4 | 0 | 2 | 8 | 16 | 6 | 40 |
| total | 12/1 | 19/5 | 45/2 | 22/2 | 31/1 | 24 | 36 | 21 | 210/11 |

12 cells not included in this table: both QC failed, 4 cells; RNA QC failed cells at blastocyst stage, 8 cells.
Red colored number indicates only RNA data passed QC.

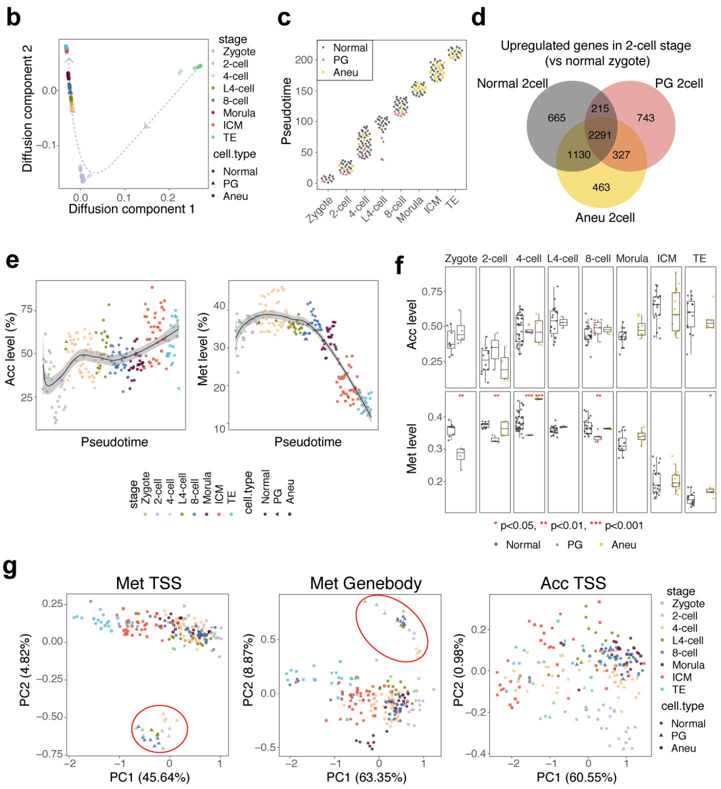


**Extended Data Figure 2 Aberrant transcriptome and epigenomes of abnormal embryos.**

(**a**) Summary of numbers of Normal, PG and Aneu cells. PG, parthenogenetic; Aneu, aneuploid.

(**b**) Developmental trajectory along preimplantation development inferred from the RNA data.

(**c**) Dot plot showing cell orders along the pseudotime (y-axis) for each stage (x-axis).

(**d**) Venn diagram showing the overlaps of 2-cell embryos upregulated genes (2-cell blastomeres vs zygotes, FDR < 0.01, fold change > 2) among Normal, PG and Aneu blastomeres.

(**e**) Dynamics of global chromatin accessibility (left) and DNA methylation level (right) of each individual cell along developmental trajectory. The black lines show the regression fit.

(**f**) Box plot showing the chromatin accessibility (top) and DNA methylation level (bottom) of Normal, PG and Aneu cells at each stage. Wilcox-test, Normal cells as control for each stage, * p < 0.05, ** p < 0.01, *** p < 0.001.

(**g**) PCA analysis of DNA methylation level at TSS (left) and gene body (middle) regions as well as chromatin accessibility at TSS (right) regions for Normal, PG and Aneu blastomeres across preimplantation development. Red cycles indicate the PG cells.

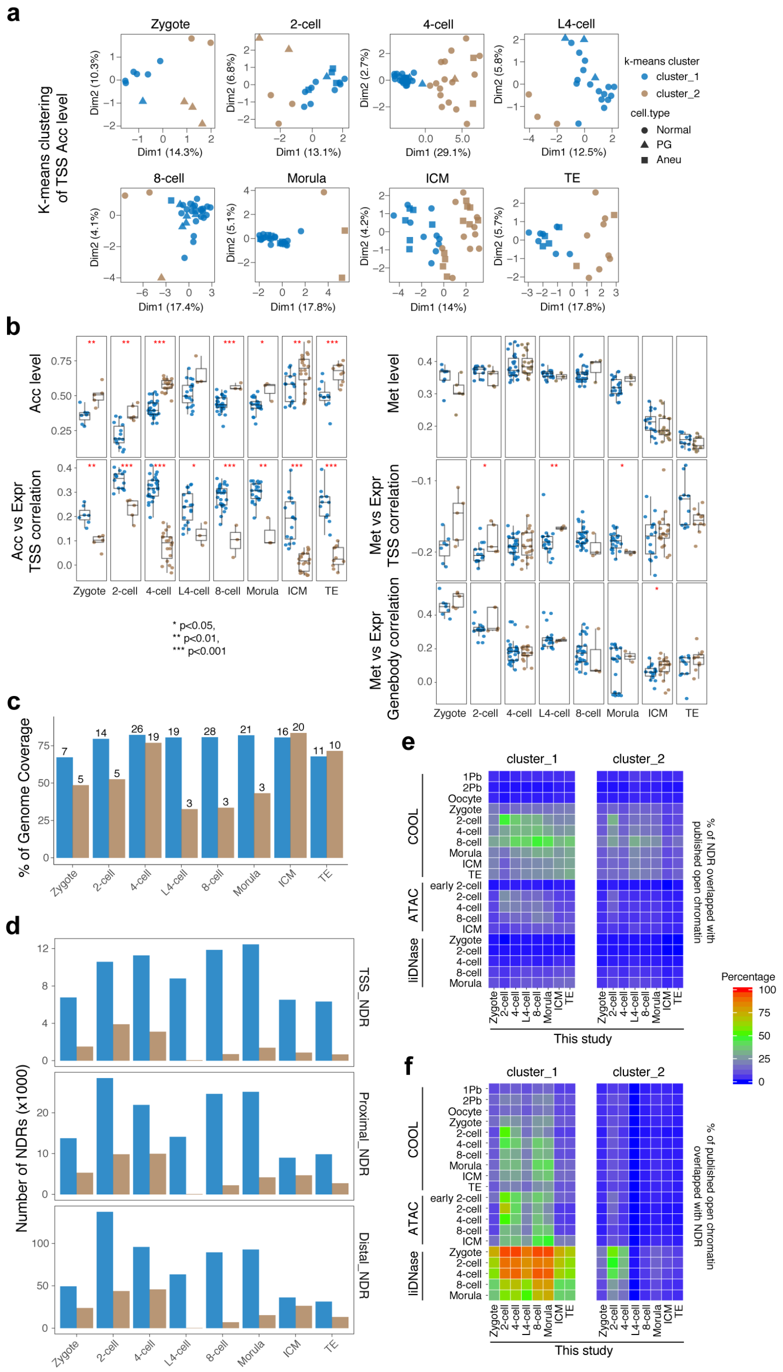


**Extended Data Figure 3 scNOMeRe-seq uncovers different subgroups with specific chromatin accessibility pattern for each stage.**

(**a**) K-means clustering (k=2) of chromatin accessibility level at TSS regions for each stage. PG, parthenogenetic; Aneu, aneuploid.

(**b**) Box plot showing the chromatin accessibility (left top), weighted Pearson correlation coefficients between chromatin accessibility at TSS regions and gene expression level (left bottom), DNA methylation level (right top), weighted Pearson correlation coefficients between DNA methylation level at TSS regions and gene expression level (right middle), weighted Pearson correlation coefficients between DNA methylation level at gene body regions and gene expression level (right bottom) for each k-means cluster at each stage. Wilcox-test, * $p < 0.05$, ** $p < 0.01$, *** $p < 0.001$.

(**c**) Bar plot showing the genomic coverage of unique GCH sites in each cluster at each stage. The number above each bar indicates the cell number of each cluster in each stage.

(**d**) Bar plot showing the numbers of TSS_NDRs, Proximal_NDRs and Distal_NDRs in each cluster at each stage.

(**e**) Heat map showing the percentages of NDRs overlapped with previously published open chromatin regions accounting for total NDRs detected in this study (ref. 4,6,10).

(**f**) Heat map showing the percentages of previously published open chromatin regions overlapped with NDRs detected in this study accounting for total published open chromatin regions (ref. 4,6,10).

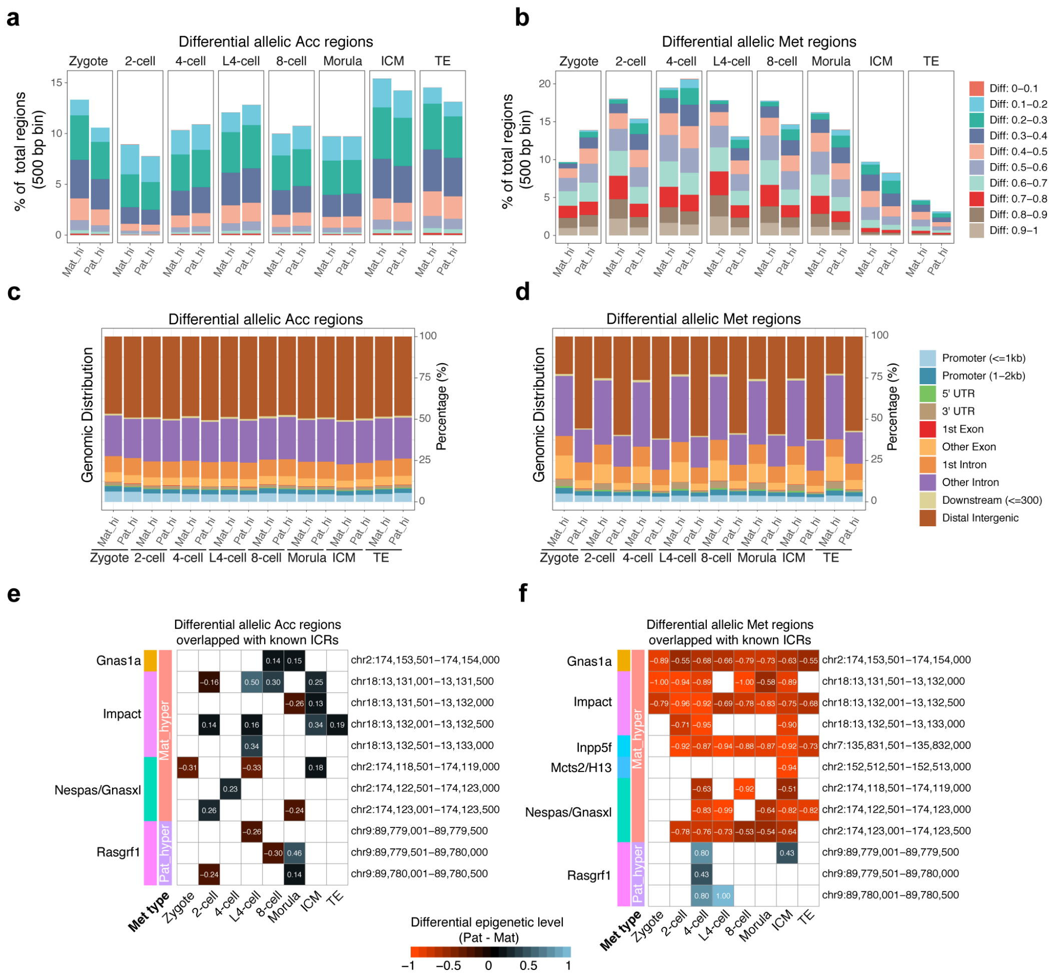


**Extended Data Figure 4 Differential allelic epigenomes in early embryos.**

(**a-b**) Bar plot showing the fractions of differential (FDR < 0.01) allelic chromatin accessibility (**a**) and DNA methylation level (**b**) between maternal genome and paternal genome of each stage. Mat_hi, regions with higher maternal allelic level; Pat_hi, regions with higher paternal allelic level.

(**c-d**) Bar plot showing the genomic distributions of differential allelic chromatin accessibility (**c**) and DNA methylation level (**d**) regions of each stage.

(**e**) Heat map showing the difference of allelic chromatin accessibility at the differential allelic chromatin accessibility regions that overlap known imprinting control regions (ICRs). Pat_hyper, known paternal hypermethylated ICRs; Mat_hyper, known maternal hypermethylated ICRs. The numbers indicate the differential chromatin accessibility level.

(**f**) Heat map showing the difference of allelic DNA methylation level at the differential allelic DNA methylation regions that overlap known ICRs. Pat_hyper, known paternal hypermethylated ICRs; Mat_hyper, known maternal hypermethylated ICRs. The numbers indicate the differential DNA methylation level.

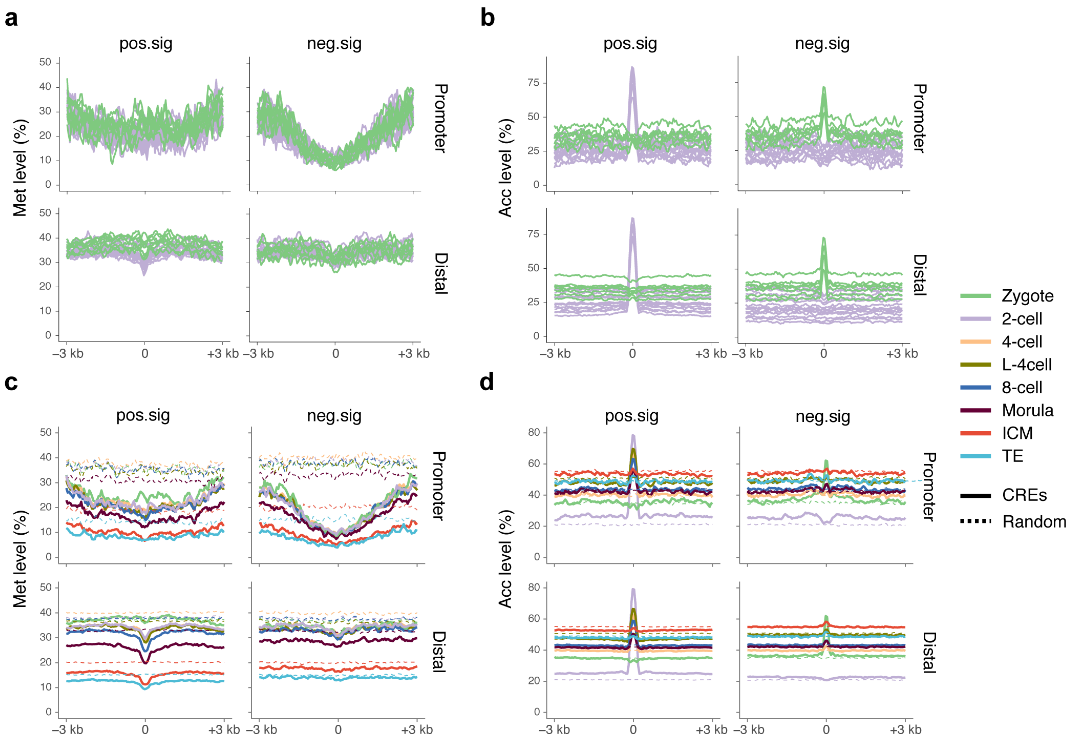


**Extended Data Figure 5 Dynamics of DNA methylation and chromatin accessibility around ZGA associated CREs.**

(**a-b**) DNA methylation level (**a**) and chromatin accessibility (**b**) around ZGA associated CREs (from the upstream 3 kb of CRE center to the downstream 3 kb) for each individual cell of zygote and 2-cell embryos.

(**c-d**) DNA methylation level (**c**) and chromatin accessibility (**d**) around ZGA associated CREs and random regions for each preimplantation stage.

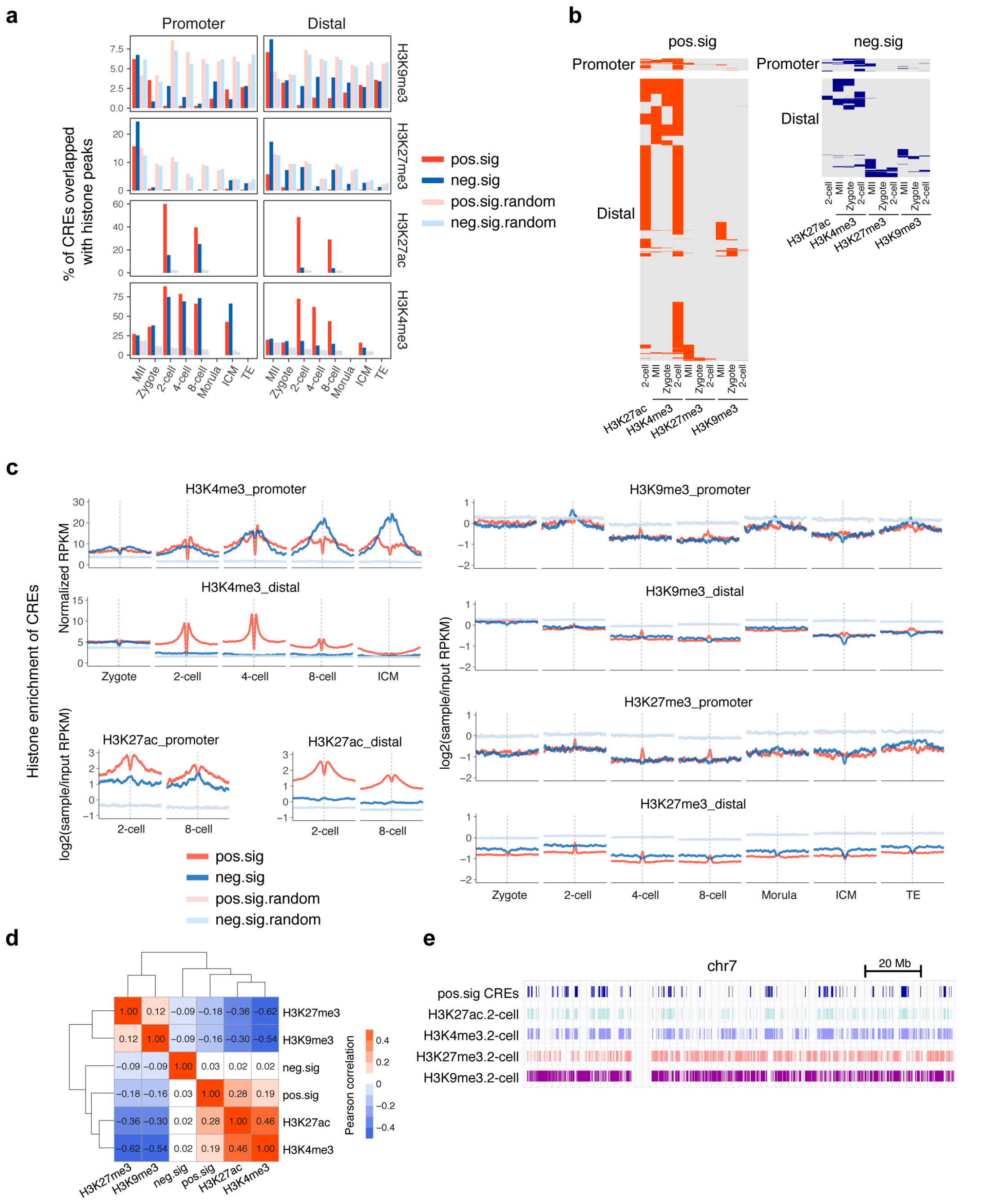


**Extended Data Figure 6 Histone modification features of ZGA associated CREs.**

(**a**) Bar plot showing the percentage of histone modification peaks overlapped with ZGA associated CREs and random regions at promoter (left) and distal (right) regions for each stage.

(**b**) Heat map showing the distribution of histone modification peaks at ZGA associated CREs.

(**c**) Profiles showing the histone modification enrichment around ZGA associated CREs and random regions for each stage (from the upstream 3 kb of region center to the downstream 3 kb).

(**d**) Heat map showing the colocalization among ZGA associated CREs and histone modification peaks of H3K27ac, H3K4me3, H3K27me3 and H3K9me3 at 2-cell stage. The numbers showing the Pearson correlation coefficients.

(**e**) IGV snapshot showing the distribution of ZGA positive-correlated CREs, histone modification peaks of H3K27ac, H3K4me3, H3K27me3 and H3K9me3 at 2-cell stage on chromosome 7.

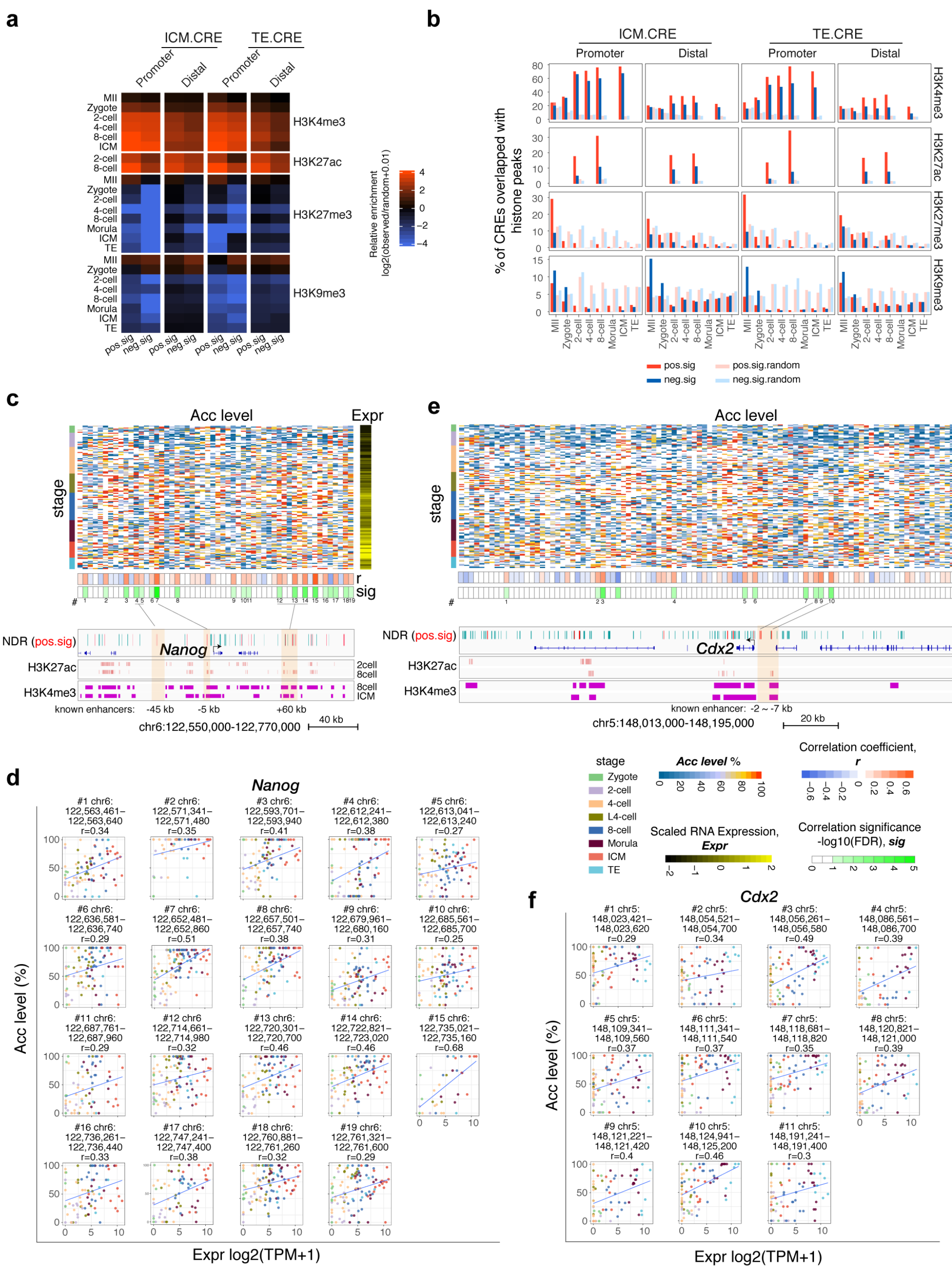

**Extended Data Figure 7 Correlation analysis reveals ICM/TE associated CREs.**

(**a**) Heat map showing the enrichment of histone modifications in ICM.CREs (left) and TE.CREs (right). The enrichment was calculated as log2 ratio for the number of observed CREs that overlap with histone modification peaks divided by the number of random regions that overlap with histone modification peaks.

(**b**) Bar plot showing the percentage of ICM.CREs (left) and TE.CREs (right) overlapped with histone modification peaks for each stage. Random regions are also shown.

(**c** and **e**) (top) Heat map showing chromatin accessibility (Acc level) of *Nanog* (**c**) or *Cdx2* (**e**) gene locus surrounding NDRs (from TSS upstream 100 kb to TES downstream 100 kb), and expression level (Expr, scaled log2(TPM+1)) of *Nanog* (**c**) or *Cdx2* (**e**) in each individual cell of early embryos; the weighted Pearson correlation coefficients (r) and significances (sig) are shown. #, the labels of positive-correlated CREs. (bottom) IGV snapshot showing the distribution of *Nanog* (**c**) or *Cdx2* (**e**) surrounding NDRs (positive-correlated CREs are in red), H3K27ac and H3K4me3 peaks. Known enhancers of *Nanog* (**c**) or *Cdx2* (**e**) are shaded.

(**d** and **f**) Scatter plot showing the expression level of *Nanog* (**d**) or *Cdx2* (**f**) and chromatin accessibility of positive-correlated CREs which have been labeled in (**c**) for *Nanog* and (**e**) for *Cdx2*. The genomic coordinates of CREs and the correlation coefficients are shown.

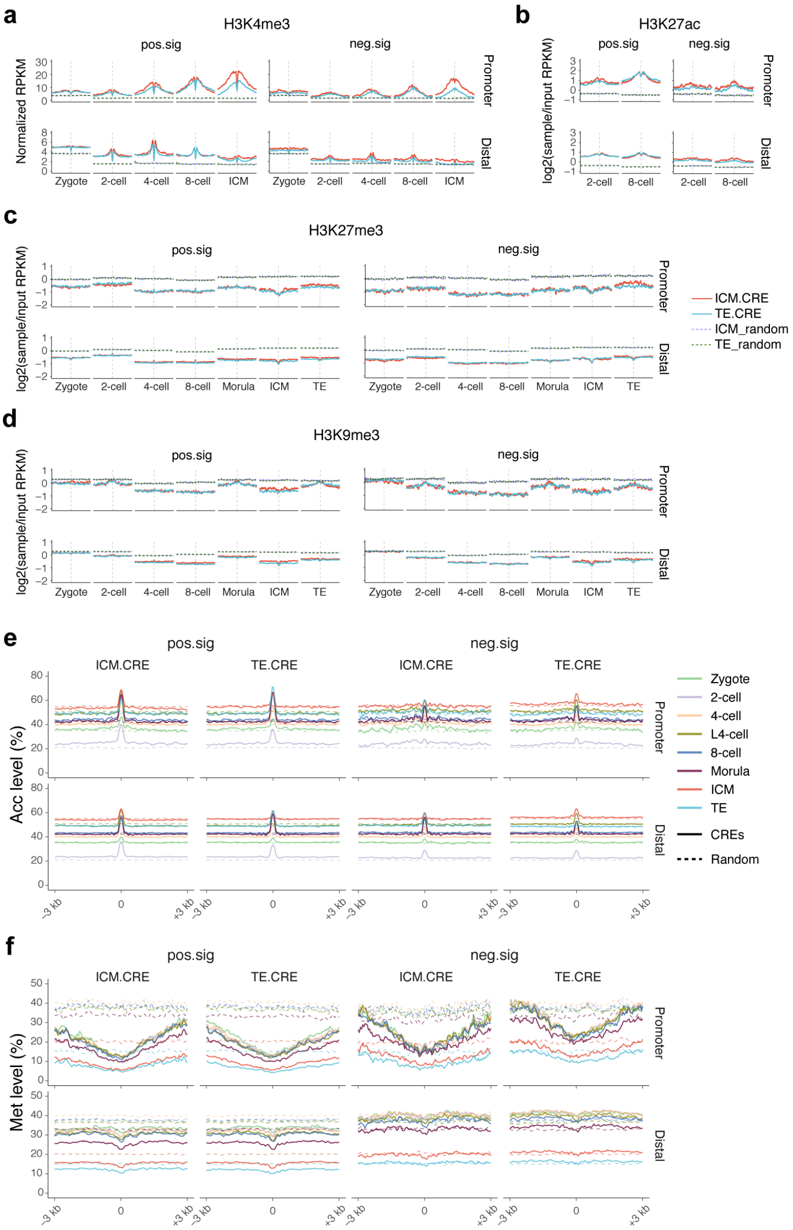


**Extended Data Figure 8 Dynamics of epigenetic level around ICM/TE associated CREs.** (**a-d**) Profiles showing the histone modification enrichment around ICM/TE associated CREs and random regions for each stage (from the upstream 3 kb of region center to the downstream 3 kb). (**e-f**) Chromatin accessibility (**e**) and DNA methylation level (**f**) around ICM/TE associated CREs and random regions for each stage.

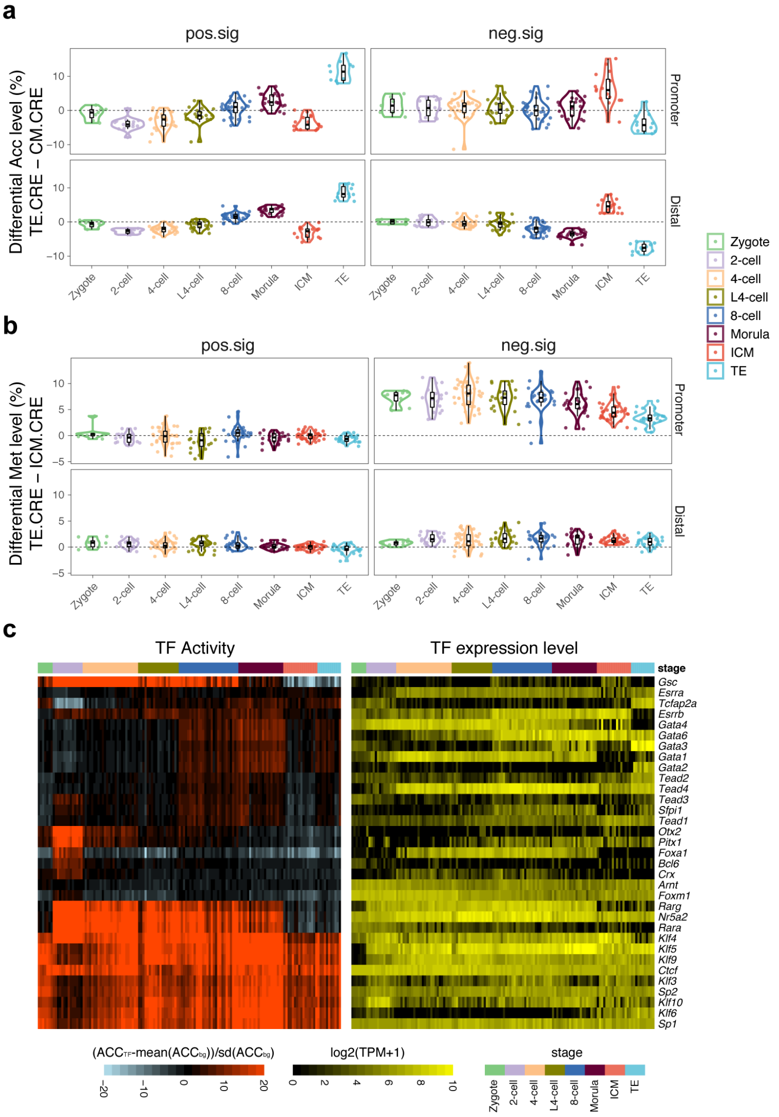


**Extended Data Figure 9 Unsynchronized activation of ICM/TE associated CREs during preimplantation development.**

(**a-b**) Violin plot showing the global difference of chromatin accessibility (**a**) and DNA methylation level (**b**) between ICM.CRE and TE.CRE in each individual cell of each stage.

(**c**) Heat map showing the TF activity (left) and RNA expression level (right) of ICM.CREs and TE.CREs enriched TFs in each individual cell across preimplantation development.
